# Metabolic glues as a means of purine sensing and chemotherapeutic response

**DOI:** 10.64898/2026.05.05.723063

**Authors:** Samuel R. Witus, Megan M. Kober, Heegwang Roh, Zhi Yang, Fouad Choueiry, Avani S. Ghate, Denis V. Titov, Michael Rapé

## Abstract

Molecular glues stabilize weak interactions to impart novel functionalities onto complexes. While plant hormones or drugs are known molecular glues, it is still unknown whether this modality provides endogenous regulation in human cells. Here, we show that purine nucleotides are molecular glues that tether the rate-limiting enzyme of purine biosynthesis, phosphoribosyl-pyrophosphate-amidotransferase (PPAT), to its inhibitor NUDT5. This mechanism allows cells to sense purine levels and establish essential feedback control of their synthesis. Thiopurine chemotherapeutics, in clinical use since the 1950’s, act as molecular glues of the same complex, but adopt unique orientations for enhanced function. Distinct from the recognition of many therapeutic glues, metabolic glue pockets can adjust their conformation to significant compound alterations and thereby enable increasing glue potency without sacrificing specificity. Our findings therefore identify endogenous metabolic glues as a mode of nutrient sensing that can be exploited to obtain compounds that rewire metabolic pathways for therapeutic benefit.

## Introduction

The ability of cells to synthesize distinct metabolites from shared precursors is essential for tissue formation and homeostasis, and the responsible biosynthetic pathways must rapidly adapt to the frequent shifts in metabolic demands that occur during proliferation, differentiation, or stress. This requires regulatory circuits that continuously monitor and adjust the levels of metabolic precursors and products ^1–4^. Loss of adequate nutrient sensing and feedback control accounts for systemic diseases, such as diabetes ^5^, or developmental pathologies as seen in patients with PRPS1 mutations or Lesch-Nyhan syndrome that stem from dysregulated purine metabolism ^6^.

Among the most critical and energetically demanding metabolic pathways, *de novo* purine biosynthesis uses ATP, amino acids, or ribose-5-phosphate to generate nucleotides that fuel proliferation, nucleic acid synthesis, signaling, and bioenergetics ^7,8^. To prevent excess consumption of metabolic building blocks, *de novo* purine synthesis is regulated by end-product inhibition ^9–13^. Conversely, high purine demand stimulates synthesis via allosteric enzyme activation, formation of a multi-enzyme purinosome, assembly of enzyme filaments, and mTORC1 or RAS-ERK signaling ^14–25^. *De novo* purine biosynthesis is complemented by a salvage pathway that recycles nucleobases at reduced energetic cost ^26^. Isotope tracing revealed that most tissues rely on both *de novo* and salvage pathways ^27^, suggesting a need for coordination that remains incompletely understood.

Reflecting its importance for cell survival, purine metabolism has been widely exploited for the treatment of cancer, autoimmune disorders, and organ transplant rejection ^28,29^. Chemotherapeutics targeting purine metabolism were introduced into the clinic in the 1940’s, including the antifolates aminopterin or methotrexate (MTX) and the thiopurine 6-mercaptopurine (6-MP) that remain staples of cancer treatment today ^30,31^. While MTX depletes metabolic cofactors required for purine synthesis ^32^, 6-MP is thought to act, at least in part, by being incorporated into DNA ^33^. Both drugs induce DNA damage to impart cancer cell death ^34^, but show off-target toxicity that limits therapeutic efficiency ^35,36^. A better understanding of purine biosynthetic pathways might not only shed light onto metabolic control circuits but also inform strategies for developing effective treatments of cancer and autoimmunity.

Here, we show that *de novo* purine synthesis is regulated by endogenous molecular glues referred to as metabolic glues. Structural studies of the rate-limiting enzyme of purine synthesis, phosphoribosyl pyrophosphate amidotransferase (PPAT), revealed that purine nucleotides anchor its inhibitor, NUDT5, in the active site of PPAT. This mechanism enables cells to monitor purine levels and provide essential feedback control for their own production. Thiopurine chemotherapeutics act as molecular glues of the same PPAT-NUDT5 complex, but adopt altered binding modes at the subunit interface that translate conformational changes in PPAT into glues with increased potency. Importantly, we could exploit the structural flexibility of this metabolic glue pocket to increase thiopurine efficiency without affecting specificity. Our study therefore identifies metabolic glues as a mode of nutrient sensing that provides starting points for the rational discovery of therapeutically relevant molecular glues.

## Results

### NUDT5 regulates *de novo* purine biosynthesis through PPAT

As Cullin-RING E3 ligases (CRLs) have emerged as critical regulators of biosynthetic pathways ^37–43^, we hypothesized that modulating their activity may allow us to discover mechanisms of metabolic control. We therefore sensitized osteosarcoma cells with sublethal doses of the CRL inhibitor MLN4924 and performed a genetic screen using an sgRNA library targeting ∼3,000 metabolic enzymes and transporters (**Fig. 1a; Extended Data Fig. 1a, b**). We found that cells depleted of enzymes of *de novo* purine synthesis, specifically those leading to adenylate production, were more sensitive to partial CRL inhibition than control cells (**Fig. 1b-d**). Conversely, cells depleted of enzymes of *de novo* pyrimidine synthesis, which compete with purine biosynthesis for shared precursors, were more resistant to CRL inhibition. Similar to targeting the pyrimidine pathway, loss of NUDT5 was protective to MLN4924 treatment (**Fig. 1b-d**). NUDT5 is a member of the Nudix hydrolase family that cleaves ADP-ribose into AMP and ribose-5-phosphate (R5P) and synthesizes nuclear ATP to facilitate chromatin remodeling and DNA repair ^44–47^, but how it regulates the response to CRL inhibition was unknown.

**Figure 1.**
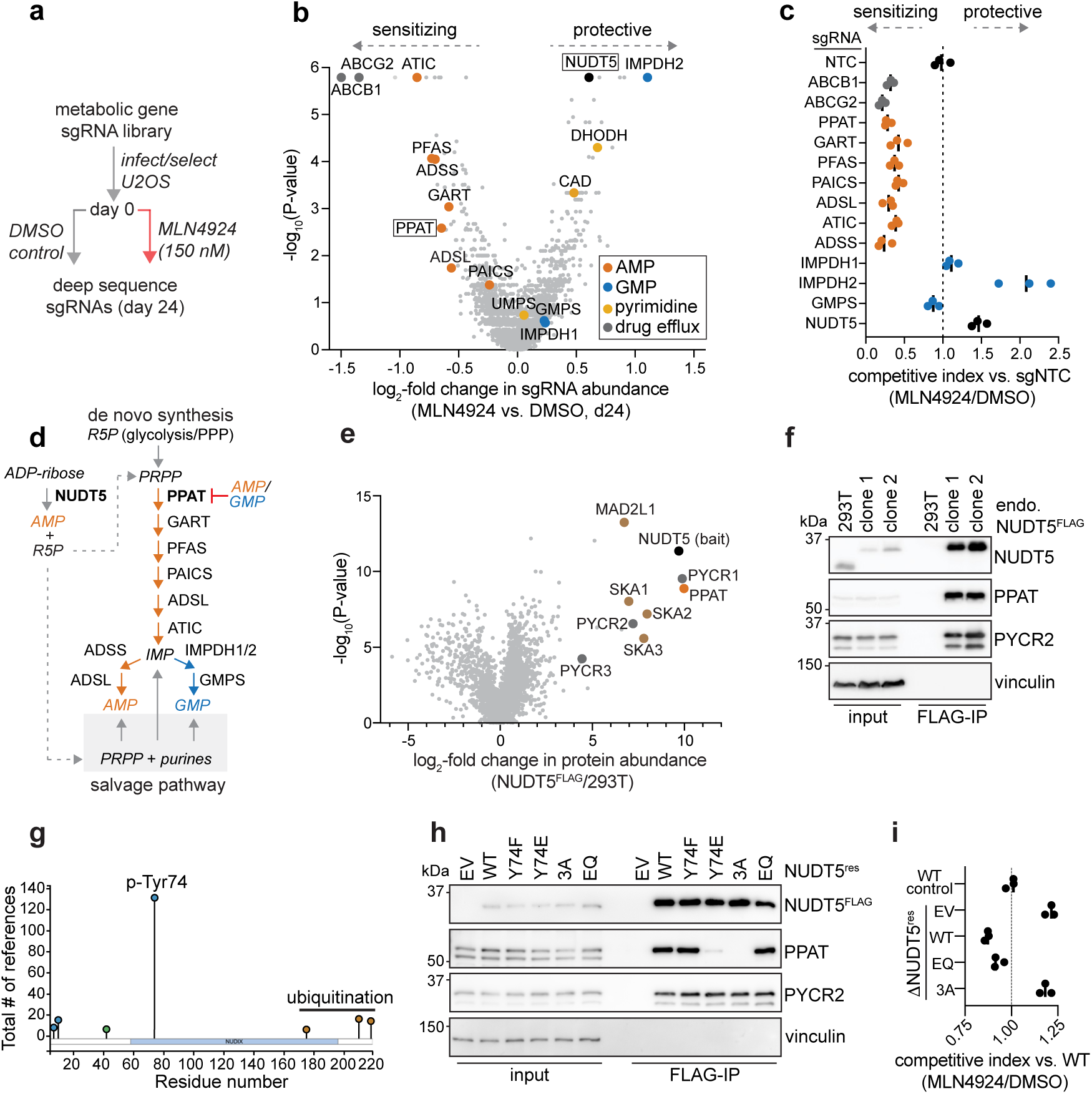
NUDT5 is a regulator of purine metabolism via interaction with PPAT. **a.** Outline of metabolism-focused CRISPR screen. **b.** Comparison of metabolic gene sgRNA abundance on day 24 between MLN4924-treated and DMSO-treated cells in CRISPR screen. **c.** Validation of MLN4924 sensitivity for select gene knockdowns via flow cytometry (FACS)-based cell competition experiment in U2OS cells. Data show n=3 biological replicates from independent experiments. **d.** Schematic of the *de novo* purine biosynthesis and purine salvage pathways in addition to the ADP-ribose hydrolase activity of NUDT5. Metabolites are italicized. **e.** Mass-spectrometry proteomics of NUDT5^3xFLAG^ immunoprecipitation from HEK293T cells. Statistical data are derived from n=3 technical replicates. **f.** Western blot of FLAG immunoprecipitation from endogenously edited NUDT5^3xFLAG^ HEK293T cells. Similar results were obtained in two independent experiments. **g.** Survey of NUDT5 post-translation modifications from the PhosphoSitePlus database filtered for n>5 observations. **h.** Western blot of FLAG immunoprecipitations from *ΔNUDT5* HEK293T cells expressing wildtype NUDT5^3xFLAG^ and mutants. Similar results were obtained in two independent experiments. **i.** FACS-based growth competition experiment comparing *ΔNUDT5* and rescue mutants to wildtype HEK293T cells treated with MLN4924. Data show n=3 biological replicates from a representative experiment. Similar results were obtained in two independent experiments.

To reveal how NUDT5’s function in MLN4924-treated cells, we identified its interactors by affinity purification and mass spectrometry. NUDT5 co-immunoprecipitated with the rate-limiting enzyme of *de novo* purine synthesis, phosphoribosyl-pyrophosphate-amidotransferase (PPAT), depletion of which sensitized cells to MLN4924 (**Fig. 1e, f**). NUDT5 also bound the mitochondrial proline biosynthetic enzymes PYCR1/2 and components of the kinetochore-associated SKA complex. Sequential immunoprecipitation of PPAT and NUDT5 showed that these proteins form a discrete complex that does not include PYCR1/2 or SKA subunits (**Extended Data Fig. 1c, d**).

AlphaFold modeling ^48^ suggested that NUDT5 recognizes PPAT through its Y74-residue that can be phosphorylated in cells (**Fig. 1g; Extended Data Fig. 2**). While a conservative Y74F mutation did not impede binding to PPAT, the phosphomimetic NUDT5^Y74E^ or a more drastic loop mutation in NUDT5^R70A/L72A/Y74A^ (NUDT5^3A^) disrupted this interaction (**Fig. 1h**), and introduction of NUDT5^3A^ into *ΔNUDT5* cells failed to restore MLN4924 sensitivity (**Fig. 1i**). By contrast, a catalytic NUDT5 mutant defective in nucleotide hydrolysis, NUDT5^E112Q/E116Q^ (NUDT5^EQ^; ref. ^44^), still bound PPAT (**Fig. 1h**) and re-established MLN4924 sensitivity in *ΔNUDT5* cells (**Fig. 1i**). These observations showed that NUDT5 controls drug sensitivity via PPAT, but independently of its enzymatic role as a Nudix-family hydrolase. Our findings are consistent with recent preprints that implicated NUDT5 as a regulator of PPAT, without having revealed a molecular basis for this activity ^49–52^.

### Structure of PPAT-NUDT5-AMP complex

To determine how NUDT5 regulates PPAT, we purified a complex between NUDT5^Y74F^ and PPAT in the presence of AMP, which had been suggested to be important for PPAT stability ^53^ (**Extended Data Fig. 3a**). The PPAT-NUDT5^Y74F^ complex was subjected to cryo-EM structure determination, yielding a density map with a global resolution of 2.8 Å that was sufficient for high-confidence model building (**Fig. 2a, b; Extended Data Fig. 3b, c, f, g; Extended Data Fig. 4a, b; Supplementary Table 1**). The structure revealed that NUDT5 dimers occupy each elliptical face of a PPAT tetramer, forming a ∼330 kDa complex. We observed density for four 4Fe-4S clusters bound by PPAT, yet did not detect its N-terminal propeptide, suggesting that PPAT is present in its mature form **(Extended Data Fig. 4c, d)**. Each NUDT5 monomer makes contacts with two PPAT protomers from opposing dimers at distinct interfaces, thereby enforcing a tetrameric form of PPAT that has been found to be more sensitive to feedback inhibition ^11,15^.

**Figure 2.**
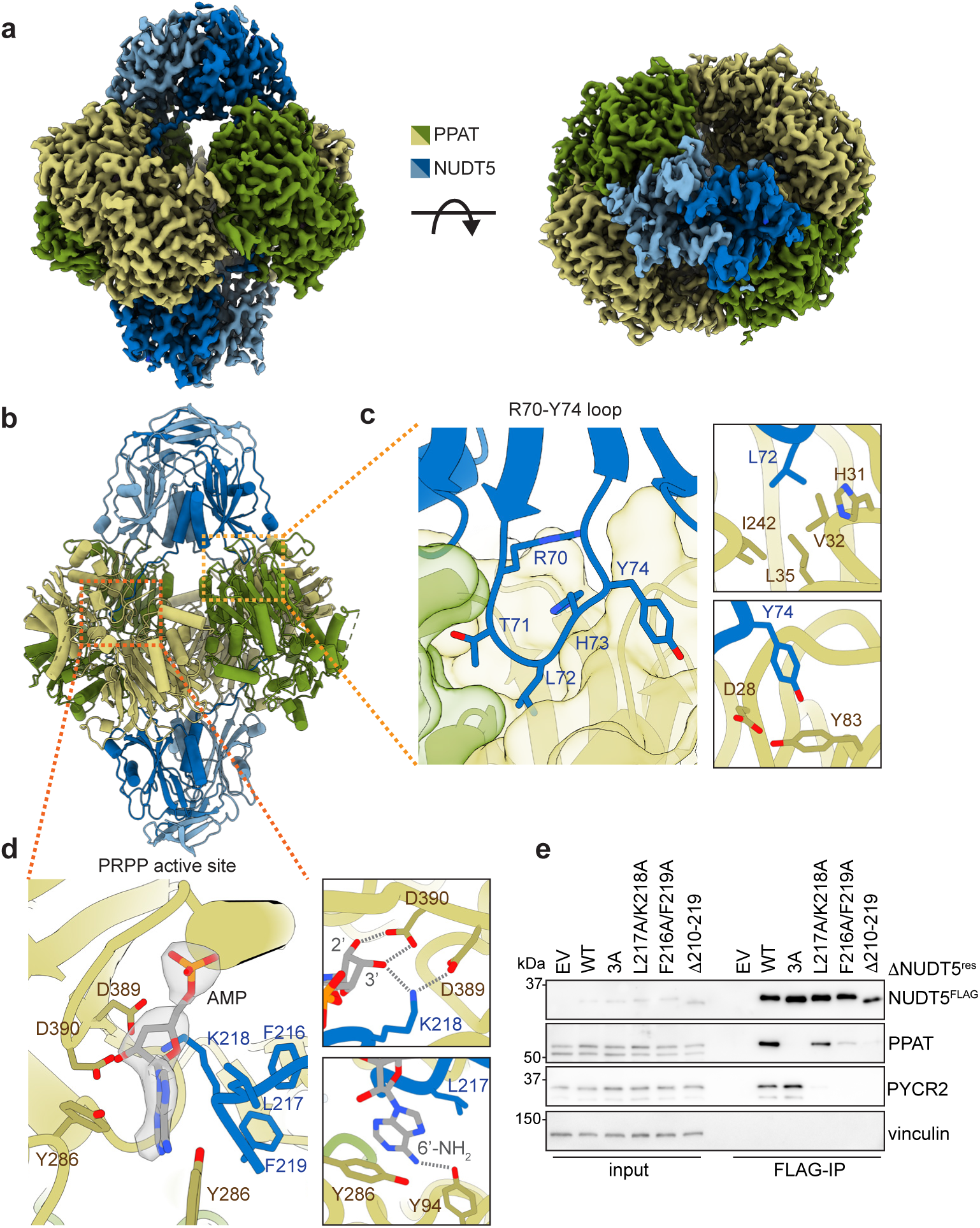
Structure of PPAT-NUDT5-AMP complex. **a.** Cryo-EM density of PPAT-NUDT5-AMP complex. **b.** Structural model of PPAT-NUDT5-AMP complex. The light orange box shows the location of the R70-Y74 loop interface, and the dark orange box indicates the positioning of the molecular glue interface from the same NUDT5 protomer. **c.** The R70-Y74 loop interface showing specific interactions between NUDT5 and PPAT including the phospho-Y74 site (right, bottom). **d.** PPAT-NUDT5-AMP molecular glue interface. Cryo-EM density for AMP is shown as a transparent surface. Specific interactions between protein components and the AMP ribose (top right) and purine ring (bottom right) are shown. **e.** Western blot of FLAG immunoprecipitations from *ΔNUDT5* HEK293T cells re-expressing wildtype NUDT5^3xFLAG^ and C-terminal tail mutants. Similar results were obtained in two independent experiments.

In agreement with our AlphaFold model, the R70-Y74 loop of NUDT5 interacts with a conserved surface of PPAT that includes a hydrophobic pocket lined by residues H31, V32, L35, and I242 **(Fig. 2c; Extended Data Fig. 4e, f)**. NUDT5-Y74 projects into a region of PPAT that includes D28, implying both repulsion and steric hindrance in a phosphorylated state. Strikingly, in a second interaction interface that was not predicted by AlphaFold, the C-terminus of NUDT5 extends into the catalytic center of PPAT, engaging a protomer in the opposing dimeric unit relative to where the R70–Y74 loop binds (**Fig. 2d; Extended Data Fig. 4g**). The NUDT5 C-terminus is anchored by an AMP molecule that occupies the PPAT active site. Reminiscent of a molecular glue stapling NUDT5 to PPAT, AMP sandwiches between PPAT and NUDT5 and contacts residues of each protein (PPAT: Y94, Y286, D390; NUDT5: L217, K218) through its purine ring and via hydrogen bonding with the ribose hydroxyls. The position of AMP in the PPAT active site is distinct from structures of bacterial PPAT orthologs which, in the absence of NUDT5, are inhibited by AMP via negative feedback ^54,55^ (**Extended Data Fig. 4h**). Deletion or mutation of conserved C-terminal NUDT5 residues that interact with AMP strongly decreased its association with PPAT (**Fig. 2e; Extended Data Fig. 4i**), which indicated that AMP may act as a molecular glue that tethers a rate-limiting metabolic enzyme, PPAT, to its inhibitor, NUDT5.

### PPAT inhibition occurs through metabolically-derived molecular glues

While plant hormones or therapeutic compounds are known molecular glues ^56–58^, it remains to be established whether endogenous metabolites can provide similar regulation in human cells. To determine if AMP is a molecular glue that inhibits purine synthesis by tethering NUDT5 to PPAT, we purified PPAT from *ΔNUDT5* cells and reconstituted its enzymatic activity **(Fig. 3a; Extended Data Fig. 5a-c)**. We then asked whether AMP impacted the ability of NUDT5 to modulate PPAT activity. High levels of NUDT5 partially inhibited PPAT in the absence of AMP, consistent with a nucleotide-independent binding site provided by the R70-Y74 loop (**Extended Data Fig. 5d**). Remarkably, the presence of AMP increased the potency of NUDT5 to shut off PPAT by ∼50-fold, thereby allowing for PPAT regulation at endogenous NUDT5 levels (**Fig. 3b**).

**Figure 3.**
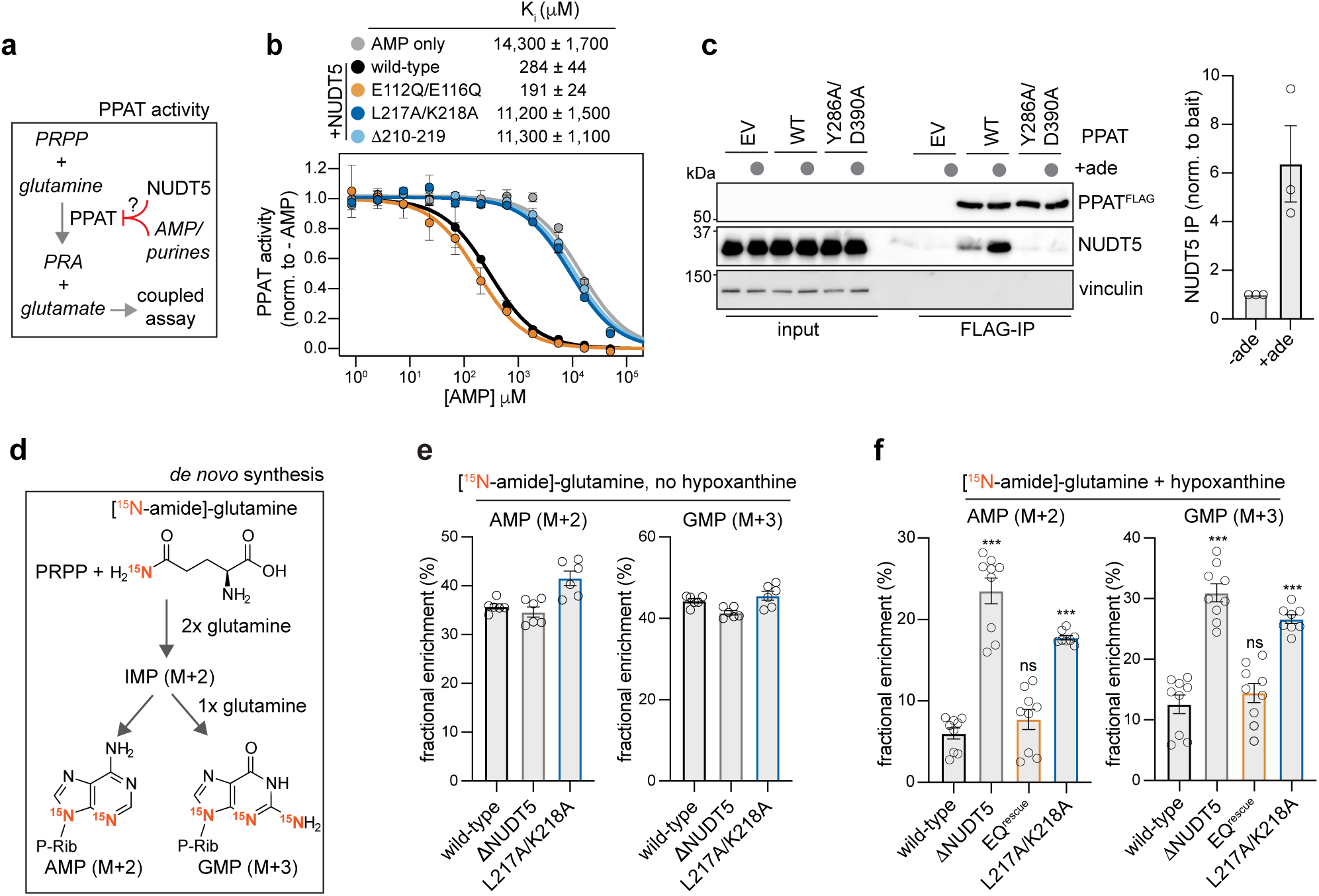
Mechanism of PPAT inhibition through NUDT5 and metabolically derived molecular glues. **a.** Biochemical activity of PPAT outlining effects of purines and NUDT5. **b.** PPAT activity assay measuring AMP-dependent inhibition in the presence and absence of wildtype NUDT5 and mutants conducted with 1 mM PRPP. Data points are the mean and error bars are SEM from n=3 independent experiments. **c.** Left – representative Western blot of FLAG immunoprecipitations from *ΔPPAT* HEK293T cells transiently expressing wildtype PPAT^3xFLAG^ or an AMP-binding deficient mutant in the presence and absence of adenine. Right – quantification of immunoprecipitated NUDT5 relative to PPAT^3xFLAG^ bait and normalized to the no adenine condition. Data show values from n=3 independent biological replicate experiments and error bars are SEM. **d.** [^15^N-amide]-glutamine labeling of purines in the *de novo* biosynthesis pathway. **e.** Fractional enrichment of AMP (M+2) and GMP (M+3) isotopologs in [^15^N-amide]-glutamine labeling experiments conducted without added hypoxanthine. Data show individual values of n=6 biological replicates from two independent experiments and error bars are SEM. **f.** Fractional enrichment of AMP (M+2) and GMP (M+3) isotopologs in [^15^N-amide]-glutamine labeling experiments conducted in the presence of hypoxanthine. “EQ” denotes NUDT5 E112Q/E116Q. Data show individual values of n=8 (L217A/K218A only) or 9 biological replicates from three independent experiments and error bars are SEM. Statistical comparisons were performed using Welch’s two-tailed t-test with Bonferroni correction between wildtype and each mutant. *** denotes a Bonferroni adjusted p-value < 0.001 and “ns” is p-value > 0.05.

To probe whether the effects of AMP onto NUDT5-inhibitor potency were mediated via a glue mechanism, we mutated the L217 and K218-residues in NUDT5 that contact AMP, but not PPAT. Importantly, both residues were required fοr AMP to stimulate NUDT5-dependent PPAT inhibition **(Fig. 3b)**. Mutation of the R70-Y74 loop that provides the second interface with PPAT also impaired NUDT5 function, in line with the notion that molecular glues stabilize weak preexisting interactions ^58^ (**Extended Data Fig. 5e**). PPAT regulation was not affected by mutations in NUDT5 that inactivate nucleotide hydrolysis **(Fig. 3b; Extended Data Fig. 5e)**. Other purines predicted to fit into the glue pocket, such as IMP, GMP, or the biosynthesis intermediate AICA-ribonucleotide, also stimulated the ability of NUDT5 to inhibit PPAT dependent on an intact glue pocket **(Extended Data Fig. 5f, g)**. Thus, multiple purine nucleotides could act as molecular glue stabilizers of an inhibited PPAT-NUDT5 complex.

**Figure 4.**
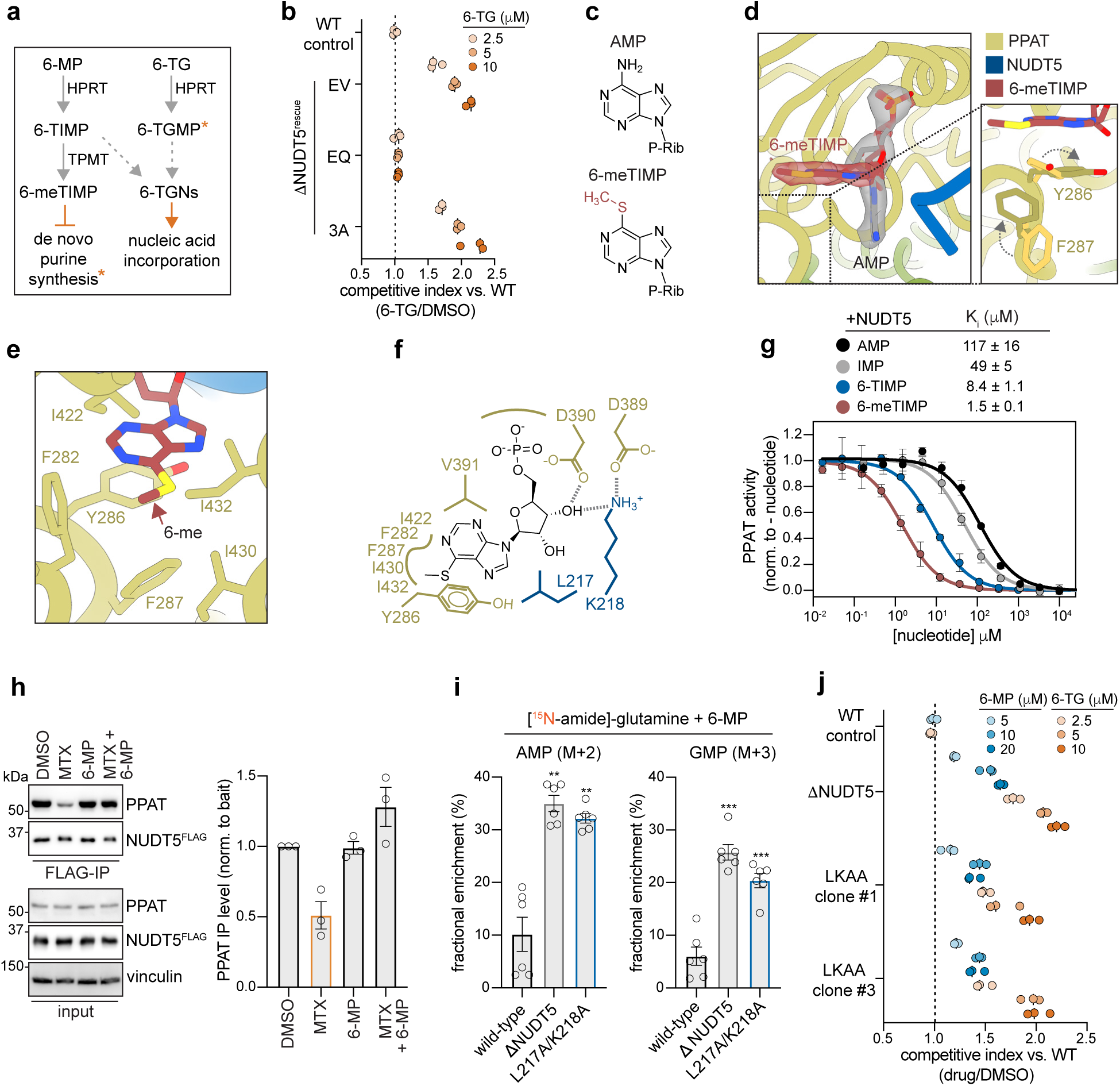
Thiopurines behave as enhanced molecular glues to inhibit PPAT. **a.** Simplified metabolism of 6-mercaptopurine (6-MP) and 6-thioguanine (6-TG). Asterisk denotes ability of 6-TGMP to be transformed into 6-meTGMP that may inhibit de novo purine synthesis. **b.** FACS-based growth competition comparing *ΔNUDT5* and mutants to wildtype HEK293T cells treated with 6-TG. Data are individual values from n=3 biological replicates from a representative experiment. Similar results were obtained in two independent experiments. **c.** Chemical structures of adenosine-5’-monophosphate (AMP) and 6-methylthioinosine-5’-monophosphate (6-meTIMP). **d.** Left – alignment of molecular glue interface of AMP and 6-meTIMP showing cryo-EM density for the nucleotides. Right – rearrangement of PPAT interface residues in the 6-meTIMP structure (dark sidechains) compared to the AMP-bounds structure (light sidechains) **e.** Hydrophobic pocket of PPAT engaged by 6-meTIMP. **f.** 2D-ligand diagram of the 6-meTIMP molecular glue interface. **g.** PPAT activity assay measuring nucleotide-dependent inhibition in the presence of NUDT5 with 0.25 mM PRPP. Data points are the mean and error bars are SEM from n=3 independent experiments. **h.** Left – Western blot of endogenous NUDT5^3xFLAG^ immunoprecipitations following 16-hour treatment with methotrexate (2 µM), 6-MP (50 µM), and MTX + 6-MP. Right – Quantification of PPAT immunoprecipitation normalized to NUDT5^3xFLAG^ bait and compared to a DMSO-treated control condition. Data are individual values from n=3 independent biological replicate experiments and error bars are SEM. **i.** Fractional enrichment of AMP (M+2) and GMP (M+3) isotopologs in [^15^N-amide]-glutamine labeling experiments conducted in the presence of 6-MP. Data points are individual values of n=6 biological replicates from two independent experiments and error bars are SEM. Statistical comparisons were performed using Welch’s two-tailed t-test with Bonferroni correction between wildtype and each mutant. *** denotes a Bonferroni adjusted p-value < 0.001 and ** is p-value < 0.01. **j.** FACS-based growth competition experiment comparing growth of *ΔNUDT5* and endogenous L217A/K218A (LKAA) NUDT5 mutants to wildtype HEK293T treated with 6-MP and 6-TG. Data show n=3 biological replicates from a representative experiment. Similar results were obtained in two independent experiments.

**Figure 5.**
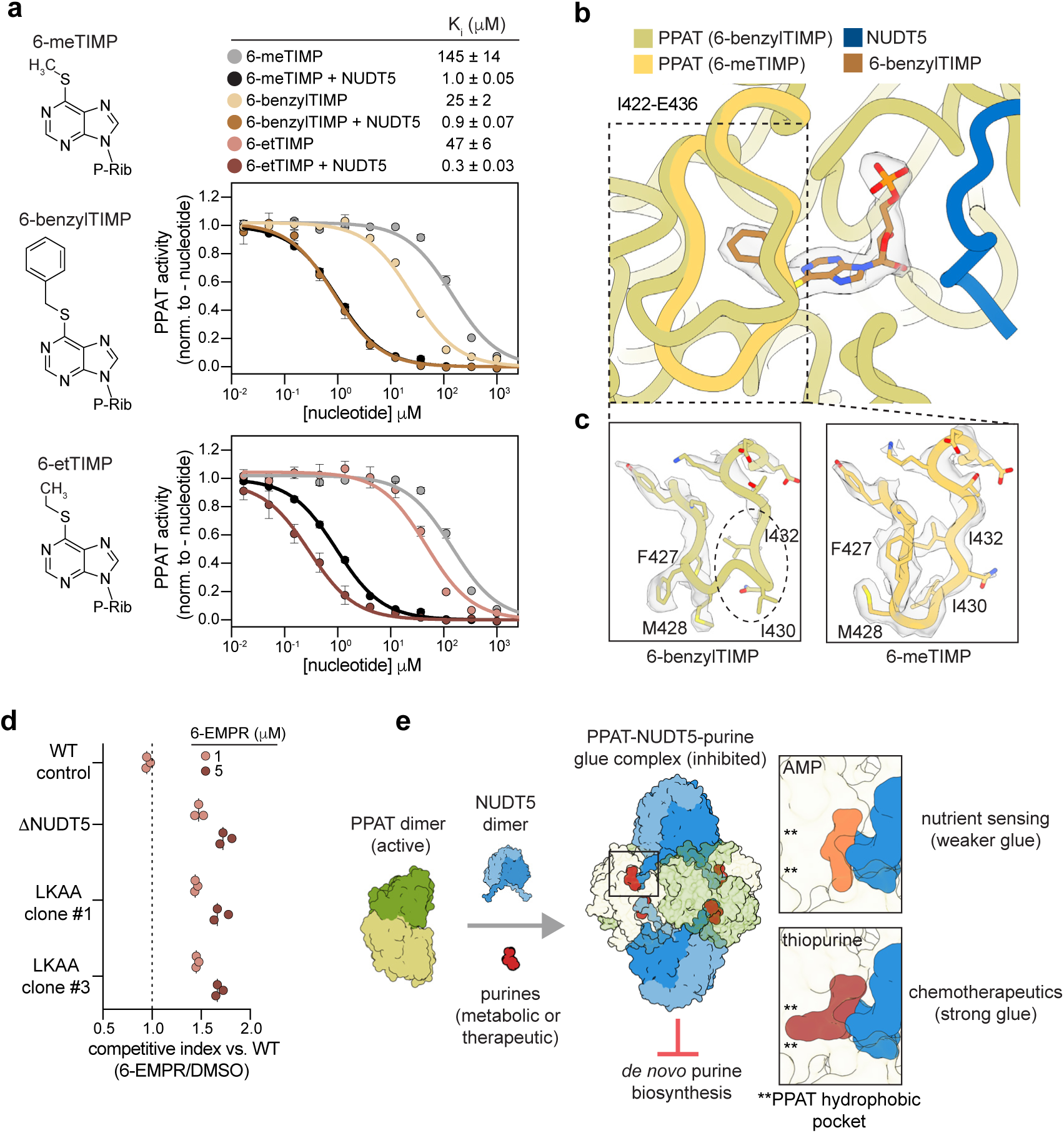
Design of improved molecular glues. **a.** Left – Molecular structures of 6-meTIMP, 6-benzylthioinosine-5’-monophosphate (6-benzylTIMP), and 6-ethylthioinosine-5’-monophosphate (6-etTIMP). Right – PPAT activity assay measuring nucleotide-dependent inhibition in the presence and absence of NUDT5. Data points are the mean and error bars are SEM from n=3 independent experiments. **b.** Structural alignment of the region surrounding the molecular glue interface for the 6-benzylTIMP- and 6-meTIMP-bound PPAT-NUDT5 structures. Cryo-EM density for the 6-benzylTIMP nucleotide is shown as a transparent surface. **c.** Sharpened cryo-EM density of the I422-E436 loop in the 6-benzylTIMP and 6-meTIMP maps shown at the same contour. The dashed oval indicates a region of missing density in the 6-benzylTIMP map that is well-defined in the 6-meTIMP map. **d.** FACS-based growth competition experiment comparing *ΔNUDT5* and endogenous NUDT5^L217A/K218A^ (LKAA) mutants to wildtype HEK293T treated with 6-ethylmercaptopurine riboside. Data show n=3 biological replicates from a representative experiment. Similar results were obtained in two independent experiments. **e.** Model of NUDT5-and purine-dependent molecular glue mechanism outlining effects on inhibition of *de novo* purine biosynthesis.

In agreement with these enzymatic studies, AMP enhanced complex formation between recombinant PPAT and NUDT5 **(Extended Data Fig. 5h)**. To assess if this regulation occurs in cells, we reconstituted *ΔPPAT* cells with either wildtype PPAT or a variant defective in AMP recognition and added adenine that is rapidly converted into AMP by the salvage pathway ^59^. Importantly, adenine treatment increased the binding of NUDT5 to PPAT, and this was blocked by mutating PPAT residues in the glue pocket (**Fig. 3c**). These data complemented our observations that deletion of the NUDT5 C-terminus abrogated complex formation (**Fig. 2e**) and further suggested that AMP behaves as a molecular glue to stabilize an inhibited PPAT-NUDT5 complex.

If stabilizing the PPAT-NUDT5 complex were relevant for signaling, loss of glue recognition should affect the ability of cells to respond to changes in purine abundance. We therefore mutated L217 and K218 in endogenous *NUDT5* to selectively ablate AMP recognition and then monitored *de novo* purine synthesis by stable isotope tracing of [^15^N-amide]-glutamine **(Fig. 3d**; **Extended Data Fig. 6a**). We conducted these experiments in the presence of hypoxanthine, which feeds the salvage pathway and should suppress *de novo* synthesis ^10^. We observed limited ^15^N-labeling of purines in either wildtype cells or cells that expressed hydrolase-inactive NUDT5^EQ^, but noted a robust increase in ^15^N-incorporation when *NUDT5* had been deleted **(Fig. 3e, f)**; this confirms that NUDT5 inhibits *de novo* synthesis in the presence of sufficient purines. Importantly, cells expressing NUDT5^L217A/K218A^ incorporated ^15^N into purines to a similar extent as *ΔNUDT5* cells (**Fig. 3f**). Total purine levels remained consistent across labeling experiments **(Extended Data Fig. 6b)**, which showed that PPAT dysregulation rather than changes in the purine pool drove enhanced *de novo* synthesis in the absence of glue recognition. These findings demonstrate that purines act as molecular glues that stabilize the PPAT-NUDT5 complex to inhibit *de novo* synthesis in the presence of sufficient purine supply. As these molecular glues are derived from endogenous metabolites, we refer to them as metabolic glues.

### Chemotherapeutics act as improved metabolic glues to inhibit purine biosynthesis

The binding of molecular glues at the interface of two proteins complicates efforts to improve the potency of therapeutic compounds ^60^, and recent work showed that chemical modification of molecular glues can in fact rewire the composition of stabilized protein complexes ^61,62^. As the PPAT-NUDT5 interface accommodates multiple purines, metabolic glue pockets may show increased flexibility and hence support recognition of small molecules with therapeutic effects. To test this notion, we turned to 6-thioguanine (6-TG) and 6-mercaptopurine (6-MP), two chemotherapeutics that are transformed in cells into metabolites that inhibit PPAT at the same site as purines ^63–65^ (**Fig. 4a**). Although these molecules were long thought to act by being incorporated in the genome to cause replicative stress, recent reports indicated that cytotoxic effects of thiopurine metabolites may be mediated, at least in part, through regulating PPAT ^49–52,66^. Indeed, we found that *ΔNUDT5* cells were highly resistant to 6-TG (**Fig. 4b**). Expression of NUDT5^3A^, which is resistant to metabolic glue regulation, did not resensitize *ΔNUDT5* cells to 6-TG, suggesting that thiopurine chemotherapeutics may act through a glue mechanism.

To test this hypothesis, we solved a 2.5 Å resolution structure of PPAT-NUDT5 bound to the 6-MP metabolite 6-methylthioinosine-5’-monophosphate (6-meTIMP) **(Extended Data Fig. 3d, f, h)**. 6-meTIMP engaged the same subunit interface between PPAT and NUDT5 as AMP **(Fig. 4c, d; Extended Data Fig. 7a)**. Surprisingly, however, the purine ring of 6-meTIMP showed a unique binding orientation in the PPAT active site compared to AMP. While 6-meTIMP and AMP are nearly aligned by their ribose-phosphates, the purine ring of 6-meTIMP adopted a bent conformation compared to the extended AMP. This is mediated by rearrangement of the PPAT Y286-and F287-residues to form a distinct π-stacking interaction between the 6-meTIMP purine ring and PPAT Y286 that is not seen in the AMP-bound structure. The re-orientation of the 6-meTIMP purine ring allows the 6-methylthio-substituent to engage a hydrophobic pocket comprised of PPAT residues F282, F287, I422, I430, and I432, which provides additional stabilizing contacts **(Fig. 4e, f)**. These findings therefore not only suggested that thiopurine metabolites act as molecular glues of the PPAT-NUDT5 complex, but they also revealed extensive flexibility of the metabolic glue pocket that endows 6-meTIMP with additional contacts to its protein partners.

Consistent with this structure, 6-meTIMP is an exceptionally strong inhibitor of PPAT that stimulated the effects of NUDT5 ∼70-fold more efficiently than AMP (**Fig. 4g**), dependent on an intact glue interface **(Extended Data Fig. 7b)**. PPAT inhibition by 6-meTIMP and NUDT5 was ∼5,000-fold more effective than by AMP alone, indicating that the combination of a therapeutic glue and its protein partner provides strict regulation of this metabolic enzyme **(Extended Data Fig. 7c)**. Addition of the thiopurine precursor of 6-meTIMP, 6-MP, to cells prevented dissociation of the PPAT-NUDT5 complex even if cells were depleted of purines, showing that enhanced glue potency enables thiopurines to overcome endogenous regulation of purine biosynthesis (**Fig. 4h; Extended Data Fig. 7d-f**). Conversely, NUDT5^L217A/K218A^-mutant cells failed to shut down purine synthesis in the presence of 6-MP, and accordingly, were highly resistant to thiopurines in growth assays (**Fig. 4i, j; Extended Data Fig. 7g**). Wildtype and mutant cells treated with 6-MP contained similar quantities of 6-TIMP and 6-meTIMP metabolites, indicating that these phenotypes are not due to a defect in purine salvage (**Extended Data Fig. 7h**). We observed similar PPAT inhibition using the methylated 6-TG metabolite 6-methylthioguanosine-5’-monophosphate (6-meTGMP), suggesting that this compound also inhibits *de novo* purine synthesis as a molecular glue (**Extended Data Fig. 7i**). We conclude that thiopurines are molecular glues that block *de novo* purine synthesis via the PPAT-NUDT5 complex. Structural flexibility in the metabolic glue pocket allows thiopurines to act more potently than endogenous nucleotides, thereby bypassing regulatory mechanisms such as allosteric PPAT activation by elevated purine precursor levels.

### Structure-based design of enhanced molecular glues

As the conformational flexibility of a metabolic glue pocket would greatly facilitate therapeutics development, we probed the consequences of attaching even bulkier substituents to 6-TIMP. We hypothesized that a 6-benzylthio group should only be accommodated if the PPAT-NUDT5 interface could undergo larger conformational changes than seen for 6-meTIMP. Intriguingly, 6-benzylTIMP retained similar PPAT inhibition as 6-meTIMP in the presence of NUDT5 and was ∼5-fold more potent in its absence **(Fig. 5a)**, indicating that the metabolic glue interface discovered here is even more pliable than anticipated.

To understand the molecular basis for flexible glue recognition, we solved the structure of 6-benzylTIMP bound to PPAT-NUDT5 (**Fig. 5b; Extended Data Fig. 3e, f, i; Extended Data Fig. 8a**). While 6-benzylTIMP binds in the same overall orientation as 6-meTIMP, the 6-benzyl group further reorients PPAT residues in the hydrophobic pocket that were observed to form stabilizing interactions with the 6-methylthio substituent of 6-meTIMP. This disrupts the local PPAT structure around a loop comprised of residues I422-E436, causing the loop to move away from the benzyl substituent and increasing flexibility in the region, as indicated by a loss of high-resolution cryo-EM density **(Fig. 5c; Extended Data Fig. 8b)**. Thus, the metabolic glue pocket in the PPAT-NUDT5 complex can undergo drastic accommodations to larger ligands in a manner that does not degrade glue potency or specificity.

Our insight into metabolic glue recognition opened the possibility to design thiopurine analogs with increased potency. We noted that the 6-methylthio substituent of 6-meTIMP occupies a hydrophobic pocket of PPAT that is not engaged by AMP. 6-meTIMP is also a more potent inhibitor than its unmethylated precursor 6-TIMP, suggesting that added hydrophobicity at this position enhances inhibitor strength **(Fig. 4g)**. In line with this notion, inclusion of an ethyl group to form 6-ethylthioinosine-5’-monophosphate (6-etTIMP) resulted in ∼3-fold greater inhibition of PPAT compared to 6-meTIMP in a manner dependent on the NUDT5 glue interface **(Fig. 5a; Extended Data Fig. 8c)**. Treatment with the prodrug, 6-ethylmercapto-purine riboside (6-EMPR) inhibited cell growth only if the glue interface was intact, supporting that cytotoxicity is mediated via NUDT5-and glue-dependent PPAT inhibition **(Fig. 5d)**. Thus, early structure-activity relationship studies reveal that the flexibility of metabolic glue pockets can be exploited to improve chemotherapeutics that had been in clinical use without alteration for many years.

## Discussion

Here, we uncovered a mechanism of regulation that is centered on a class of molecules referred to as metabolic glues (**Fig. 5e**). Metabolic glues restrict *de novo* purine synthesis under nutrient-replete conditions, showing that the glue mechanism provides a means of purine sensing to safeguard cells from undue consumption of metabolic precursors. Our work therefore establishes that endogenous molecular glues serve crucial regulatory functions in human cells. Building upon this finding, we showed that thiopurine chemotherapeutics, clinically used for ∼70 years, act as enhanced metabolic glues to inhibit *de novo* purine synthesis and suppress cell growth. Structural comparison of multiple drug-bound complexes revealed striking conformational flexibility of the metabolic glue pocket, indicating that investigation of metabolic control mechanisms may provide a fruitful avenue to discover therapeutically relevant molecular glues.

Foundational work had shown that molecular glues enhance weak, preexisting interactions to bring affinities between binding partners into a biologically meaningful range ^67–72^. Consistent with this principle, each NUDT5 protomer engages PPAT at two distinct regions to stabilize the inhibited complex, only one of which is stabilized by the molecular glue. At the glue interface, the NUDT5 C-terminal tail is anchored at the PPAT active site through its interactions with the nucleotide and surrounding regions of PPAT. This interaction is critical for PPAT feedback inhibition, allowing cells to detect purines at micromolar levels that reflect endogenous purine mono-phosphate concentrations of 10-40 μM ^73^. We conclude that metabolic glues provide a means for purine sensing that helps cells prevent excessive use of metabolic building blocks.

Although metabolic glues serve as a primary mode of regulation, they act by reinforcing a second site centered on the R70-Y74 loop of NUDT5. The R70-Y74 interface is likely broken by phosphorylation of Y74, which could couple complex formation to signaling inputs from growth factors or the cell cycle. Combined regulation by metabolic glues and phosphorylation may allow cells to quickly upregulate purine production during replication, DNA repair, or upon transcriptional bursting, which require increased levels of purine nucleotides ^74^. We expect that metabolic glues frequently cooperate with additional forms of regulation to rapidly adjust biosynthetic pathways to the complex needs of a eukaryotic cell. Metabolic glues could therefore provide a mechanism to embed biosynthetic control within larger signaling networks.

Our discovery of glue-dependent regulation of purine synthesis provides practical insight to overcome limitations in glue discovery. Unlike previously studied therapeutic glues that bind rigid and narrowly-defined pockets, the conformational flexibility of the metabolic glue pocket permits substantial ligand modifications without sacrificing inhibitor potency or specificity. We predict that such flexibility is a general feature of metabolic glue pockets, reflecting evolutionary adaptation of biosynthetic enzymes to diverse substrates and metabolic regulators. As new proteomic platforms enable identification of weak protein-metabolite interactions and subsequent discovery of glue-regulated complexes ^75^, leveraging the flexibility of metabolic glue pockets should enable rational development of therapeutic glues that are inspired by endogenous regulation. We therefore anticipate that the study of metabolic glues will not only provide more examples of nutrient sensing, but also expand the therapeutic potential of molecular glues as an emerging drug class.

## Methods

### Data reporting

No statistical methods were used to predetermine sample size. The experiments were not randomized and the investigators were not blinded to allocation during experiments and outcome assessment.

### Mammalian Cell Culture

U2OS and HEK293T cells were maintained in DMEM with Glutamax (Gibco, 10566016) supplemented with 10% FBS (VWR, 89510-186) and 1x penicillin-streptomycin (Pen-Strep; Gibco, 15140122) at 37 °C and 5% CO2. Growth media for PPAT knockout cells was supplemented with 100 µM adenine. For drug-treatment immunoprecipitation experiments, cells were plated in DMEM with Glutamax, 10% dialyzed FBS (dFBS; Cytiva, SH30079.02), and 1x Pen-Strep. For metabolomics experiments, cells were plated in DMEM (high glucose, no glutamine; Gibco, 11960044) supplemented with 2 mM L-glutamine (Gibco, A2916801), 10% dFBS, and 1x Pen-Strep. Additional details about cell culture conditions for isotopic tracing experiments can be found under “Mass-spectrometry based metabolomics”. Expi293F cells were maintained in Epxi293 media (Gibco; A1435101) in shaker flasks at 130 rpm and 8% CO2. Cell stocks were obtained from and authenticated by the UC Berkeley Cell Culture Facility and were routinely tested for mycoplasma contamination.

### CRISPR editing and endogenous cell line generation

To generate CRISPR knockout NUDT5 and PPAT lines in HEK293T and Expi293F cells, we utilized a mixture of three optimized sgRNA’s (Gene Knockout Kit V2, Synthego). 1.5 μL of sgRNA mixture (100 μM) was incubated with 3.2 μL of Cas9-NLS protein (40 μM; produced by UC Berkeley MacroLab) for 10 minutes at room temperature. RNPs were nucleofected into 5x10^5^ cells resuspended in Lonza SF solution (Lonza, V4XC-2032) using a Lonza 4D Nucleofector X-unit and pulse code CM-130. Cells were recovered in 6-well plates for 3-4 days, split once, and then cloned by single-cell dilution in 96-well plates. Expi293F cells were not cloned, as NUDT5 bulk depletion was observed to be >95% by Western blot. Clones were assessed for knockout by Western blotting and phenotypic assay. To establish a HEK293T NUDT5 knockout line used for experiments, five clones were combined at equal cell number into one pool to minimize clonal bias effects. A similar editing strategy was used for endogenous 3xFLAG epitope tagging and NUDT5 L217A/K218A mutant generation, except that a single sgRNA was used and 1 μL of Alt-R HDR donor ssODN donor template (100 μM stock; IDT) was included in the nucleofection mixture added after RNP formation. For knock-in generation, cells were recovered in 1 μM HDR Enhancer V2 (IDT) for 16 hours post nucleofection followed by a media change. Knock-in clones were validated by sanger sequencing. sgRNA and ssODN sequences used are listed below:

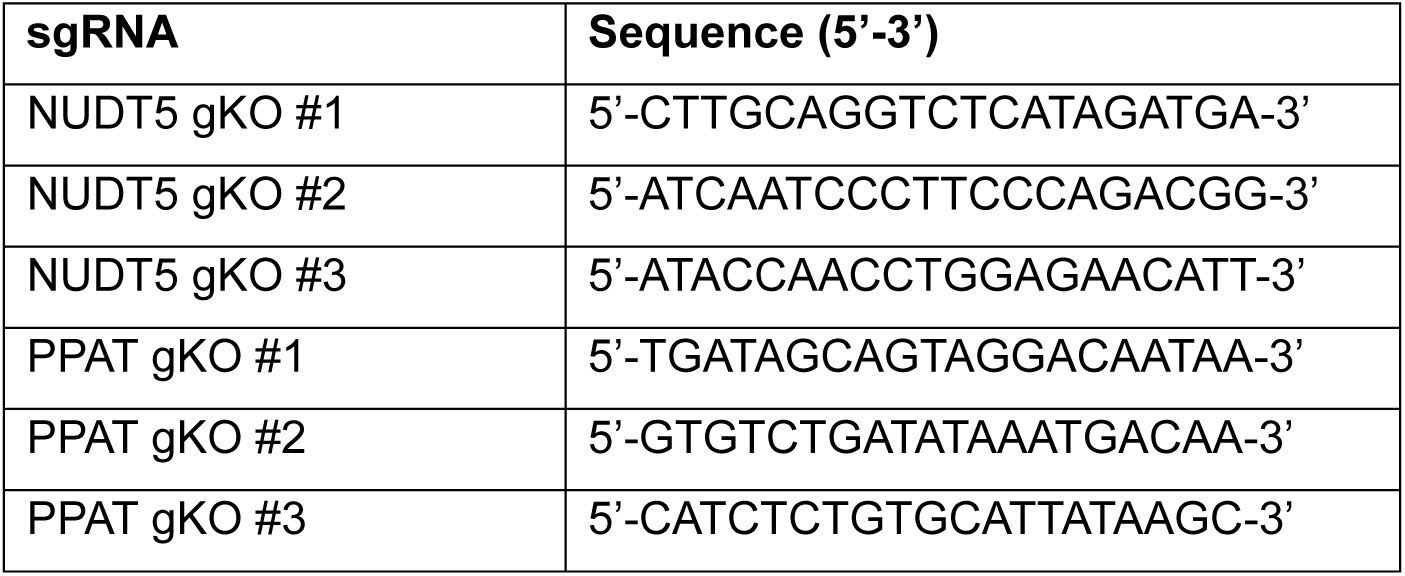
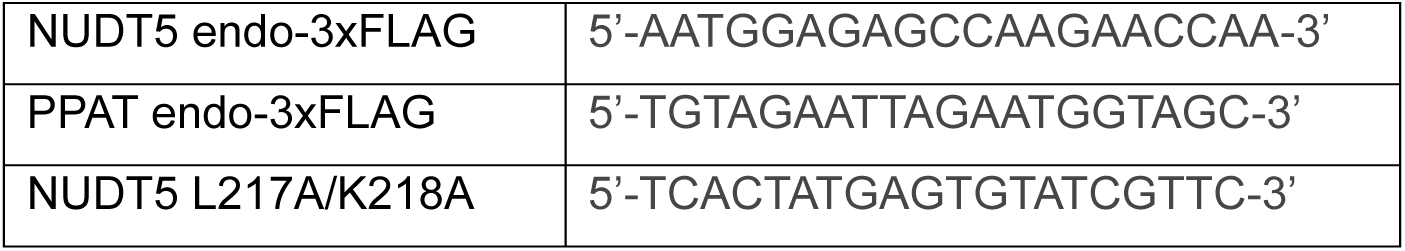

ssODN sequences:

3xFLAG-NUDT5:5’-AACTTCTCACCTGAGGGCTGTAAAGACTCGTTTGAAAATGGACTACAAGGACCACGACGGCGATTATAAGGATCACGACATCGACTACAAAGACGACGATGACAAGGGTGGGAGTGGCGGGAGTGAGAGCCAAGAACCAACGGAATCTTCTCAGAATGGCAAACAGTATATCATTTC-3’

PPAT-3xFLAG:5’-AGCTTGTCTCACTGGAAAATATCCTGTAGAATTAGAATGGGGTGGCAGCGGCGGTTCTGGTGGCAGCGACTACAAGGACCACGACGGCGATTATAAGGATCACGACATCGACTACAAAGACGACGATGACAAGTAGCTGGTAGGGTTGGATGTGTGTAGTTTCAAGATAGAAAG-3’

NUDT5L217A/K218A:5’-TCGTTTACAAAAATGGCCAGTGTCATATTTGGGCTTAAAATGCAGCGAAGGGCACTTCAAATGGCTTTGCATTTGCATGTTTCAGT-3’

### Stable cell line generation

Lentiviral particles were generated by co-transfecting pLVX-IRES-puromycin or blasticidin constructs encoding desired protein mutants with 2^nd^ generation lentiviral packaging plasmids into HEK293T cells using Mirus Trans-IT LT1 transfection reagent (Mirus, MIR 2304). For FACS-based growth competition experiments, pLVX-mCherry-P2A-blasticidin and pLVX-GFP-P2A-blasticidin constructs were used to establish fluorescent cell lines for co-culture. Following transfection of the plasmid mixture, media was changed the next day, and virus was harvested 48 hours after the media change by filtering through a 0.45 µM filter. For stable cell line generation, cells were infected with lentiviral particles at an MOI < 1 in media supplemented with polybrene (8 μg/mL). Cells were selected with antibiotic (1 μg/mL puromycin or 10 μg/mL blasticidin) ∼48 hours after transduction until all cells on a control plate were dead, at which point selected cells were recovered without antibiotics for at least 2 days prior to assaying.

### Metabolism-focused CRISPR screen

For the MLN4924 metabolism-focused CRISPR screen, we used a custom sgRNA library with 8 sgRNAs per gene targeting ∼3,000 metabolic enzymes and transporters cloned into the pLentiCRISPR-V2 vector (gift of Kivanç Birsoy, Rockefeller University). Lentiviral particles were generated by co-transfecting the sgRNA library with 3^rd^ generation lentiviral packaging plasmids into HEK293T cells using Mirus Trans-IT LT1 transfection reagent (Mirus, MIR 2304). U2OS cells grown in DMEM with Glutamax, 10% FBS, and 1x Pen-Strep were transduced with lentivirus at an MOI ∼0.5 and 500-fold sgRNA coverage in media supplemented with 8 μg/mL polybrene for 48 hours. Transduced cells were selected with puromycin (1.5 μg/mL) for 4 days, and recovered for an additional 2 days, after which a zero time-point was harvested for the screen (30x10^6^ cells). The selected and expanded cells were split into two pools maintaining 1,000-fold sgRNA coverage throughout the screen and treated with either treated with DMSO or sub-lethal doses of MLN4924 (150 nM). Cells were split every 4 days, and fresh DMSO or MLN4924 was added after allowing cells to re-adhere to plates for 4 hours. On day 24, 30x10^6^ cells were harvested per condition and genomic DNA was extracted using a QIAamp DNA Blood Maxi kit (Qiagen, 51192) according to the manufacturer’s protocols. sgRNA-encoding regions were amplified with ExTaq DNA polymerase (Takara, RR001) with unique barcoded primers. PCR products were combined and 50 μL was gel purified from a 3% agarose TBE gel. Amplicons were sequenced using an Illumina NextSeq2000 by UC Berkeley QB3 Genomics core facility. Data were analyzed using the MAGeCK analysis package ^76^, comparing normalized sgRNA representation on day 24 between DMSO and MLN4924 treated cell pools.

For screen validation, competition experiments were conducted as outlined in the “Cellular growth assays” methods section, with the following sgRNA sequences that were not present in the initial metabolism-focused screening library:

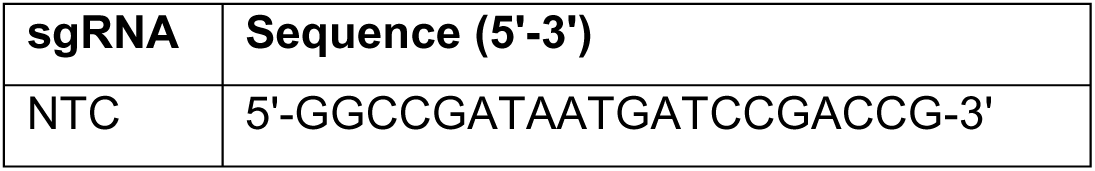
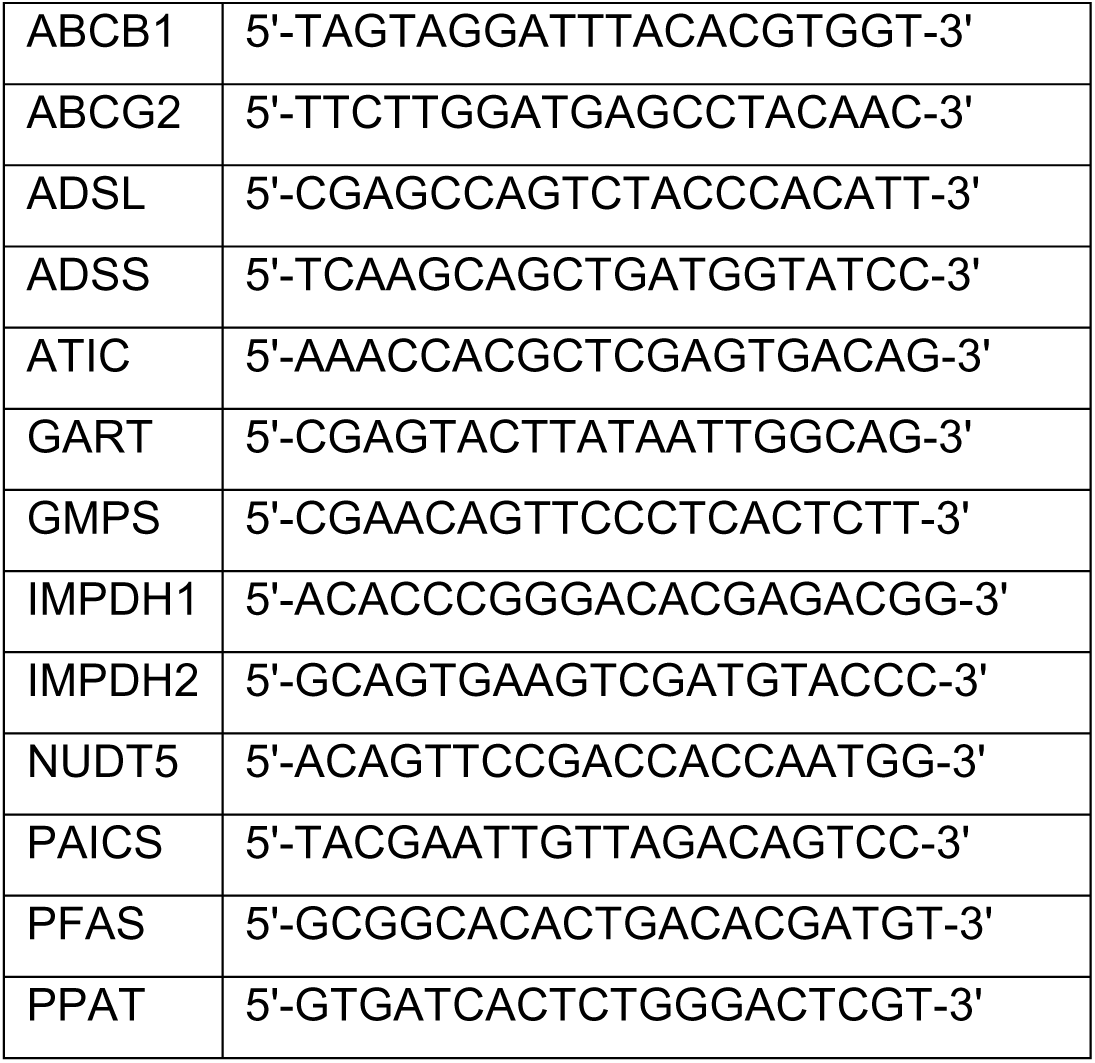

### Cellular growth assays

All growth assays were conducted in DMEM with Glutamax supplemented with 10% FBS and 1x Pen-Strep. Specific drugs are noted under “Drugs and chemicals” and amounts are specified in figures and legends.

#### FACS-based growth competition

For growth competition experiments, GFP- or mCherry-expressing cell populations for comparison were established using lentiviral expression as described above. For CRISPR screening validation experiments, GFP-expressing U2OS cells were transduced with pLentiCRISPR-V2 lentivirus particles containing sgRNA’s targeting the indicated genes (sgRNA sequences listed above) and an mCherry population was transduced with a non-targeting control sgRNA (sgNTC). After puromycin selection and recovery, GFP-knockout and mCherry-sgNTC cells were mixed in 12-well plates in equal cell number, and either DMSO or MLN4924 (100 nM) was added. Cells were split every four days, re-adding drug at each split. After 12 days, the ratio of GFP^+^ to mCherry^+^ cells was determined by flow cytometry using a BD LSRFortessa instrument and analyzed using FlowJo (v10.8.1). GFP^+^ to mCherry^+^ cell ratios for each knockdown competition condition in the presence of MLN4924 were normalized to equivalent DMSO-treated control conditions. The FACS gating strategy for growth competition experiments is outlined in Supplementary Figure 1.

For drug treatment growth competition assays using HEK293T cells, GFP^+^ and mCherry^+^ cell populations were established as described above. For rescue experiments, NUDT5 was reintroduced to *ΔNUDT5* cells via lentiviral expression using a pLVX-^3xFLAG^NUDT5-IRES-puromycin construct as described above. Briefly, 2.5x10^4^ wild-type HEK293T mCherry^+^ cells were co-plated with 2.5x10^4^ GFP^+^ cells in 12-well plates. Drug was added the following day, and cells were analyzed by flow cytometry 72 hours after addition. GFP^+^ to mCherry^+^ cell ratios for drug treatment wells were normalized to equivalent DMSO-treated cell mixtures.

#### CellTiter Glo

2.5x10^3^ U2OS cells were seeded in 96-well clear bottom plates (Corning, 165306) in 50 μL of media. MLN4924 was added the following day at 2x concentration in 50 μL of media. Viability was measured after 72 hours by addition of 100 μL CellTiter Glo reagent per well (Promega, G7570) according to manufacturer’s instructions and a luminescence signal measured on a Perkin Elmer EnVision microplate reader.

#### Incucyte

5x10^3^ HEK293T cells were seeded in black 96-well clear bottom plates in in 100 μL of media (Corning, #3904). Drugs were added the following day at 2x concentration in 100 μL of media. Cell growth was monitored using time-resolved microscopy in an Incucyte S3 microscope.

### Drugs and chemicals

The following drugs and chemicals were used in this study at amounts specified in figures and legends: Pevonedistat; MLN4924 (MedChemExpress, HY-70062), methotrexate; MTX (MedChemExpress, HY-14519), lometrexol; LMX (MedChemExpress, HY-14521), brequinar (MedChemExpress, HY-108325) , rapamycin (Adooq Biosciences, A10782), 5-Phospho-D-ribose 1-diphosphate; PRPP (Sigma-Aldrich, P8296) , L-glutamine (Sigma-Aldrich, G8540); adenosine-5’-monophosphate; AMP (Sigma-Aldrich 01930), inosine-5’-monophosphate; IMP (MedChemExpress, HY-W010759), guanosine-5’-monophosphate; GMP (Sigma-Aldrich, G8377), AICA-ribonucleotide (Cayman Chemicals 33907), adenine (Thermo Scientific, A17622.14), hypoxanthine (MedChemExpress, HY-N0091), 6-thioguanine; 6-TG (Thermo Scientific, B21280.03), 6-mercaptopurine; 6-MP (Adooq Biosciences, A15898), 6-thioinosine-5’-monophosphate; 6-TIMP (Jena Biosciences, NU-1148), 6-methylthioinosine-5’-monophosphate; 6-meTIMP (Jena Biosciences, NU-1226), 6-methylthioguanosine-5’-monophosphate; 6-meTGMP (Jena Biosciences, NU-1128), 6-benzylthioinosine-5’-monophosphate; 6-benzylTIMP (WuXi, custom synthesis), 6-ethylthioinosine-5’-monophosphate 6-etTIMP (WuXi, custom synthesis), 6-ethylmercaptopurine riboside; 6-EMPR (WuXi, custom synthesis). LC-MS and HNMR/CNMR spectra for chemicals synthesized by WuXi are attached as Supplemental Figure 2.

### Immunoprecipitations

#### Small-scale immunoprecipitations

Cells were collected from one 10 cm plate per condition by aspirating media, scraping and washing in cold DPBS, and pellets were flash frozen in liquid nitrogen or immediately processed. Cell pellets were lysed in in 500 μL lysis buffer (30 mM HEPES-NaOH pH 7.5, 150 mM NaCl, 5 mM MgCl_2_, 0.1% NP40) supplemented with EDTA-free protease inhibitors (Roche, 11873580001), PhosStop phosphatase inhibitors (Roche, 4906845001), and benzonase (Millipore, 70746). Lysis was allowed to proceed for 30 minutes with rotation, and lysates were clarified by centrifugation at 21,000 x g. An input sample was taken from the supernatant by mixing 1:1 with 2x urea sample buffer (120 mM Tris pH 6.8, 4% SDS, 4 M urea, 20% glycerol and bromophenol blue), and the remaining supernatant was incubated with FLAG-M2 agarose beads (15 μL of 50% slurry/condition; Millipore, A2220) for 2 hours at 4 °C with rotation. Beads were then washed three times with IP buffer and eluted at 65 °C with 1x urea sample buffer.

For transient expression PPAT^3xFLAG^ IP’s, *ΔPPAT* HEK293T cells were seeded at a density of 2x10^6^ cells per 10-cm dish in complete growth medium supplemented with 100 μL adenine. After 48 hours, cells were transfected with 1 μg of plasmid DNA using TransIT-293 (Mirus, MIR 2700) transfection reagent. The following day, the culture medium was replaced with adenine-free medium containing dFBS after a brief rinse with adenine-free medium. 16 hours after adenine withdrawal, adenine was re-added to a final concentration of 100 μM, and cells were harvested 4 hours later for immunoprecipitation as described above.

#### Large-scale immunoprecipitations for mass-spectrometry (IP-MS) proteomics

For IP-MS, cells were harvested from 10x15 cm plates per condition, lysed in 5 mL of lysis buffer, and processed as described above for the NUDT5^3xFLAG^ IP using 90 µL of FLAG-M2 resin slurry per sample. For the PPAT^V5-TwinStrep^ and NUDT5^3xFLAG^ sequential IP, supernatant was first bound to Streptactin XT 4Flow resin (IBA, 2-5010-010), washed 3x in lysis buffer, and eluted in lysis buffer containing 50 mM biotin (2-1016-005). Eluate was then incubated with FLAG-M2 affinity resin for two hours, followed by 3x washes in lysis buffer. An additional three DPBS washes were performed after washes in lysis buffer to remove detergent. Control immunoprecipitation from wild-type HEK293T cells was performed in parallel for all IP-MS experiments. FLAG-M2 beads were flash frozen and further processed by the UC San Diego Proteomics Facility as described below under “Mass spectrometry-based proteomics.”

#### In vitro PPAT^Strep^ pull-down

Recombinant PPAT^V5-TwinStrep^ and NUDT5^HA^ proteins were diluted in lysis buffer either supplemented with 2 mM AMP or water. For each reaction, 100 μL of 100 nM PPAT solution was mixed with 100 μL of NUDT5 at concentrations ranging from 10 μM to 2.4 nM (4-fold serial dilution). The mixtures were incubated for 1 hour at room temperature on a rotator, followed by addition of 5 μL slurry of StrepTactin XT magnetic beads (IBA, 2-5090-002) and incubated for 1 hour at 4 °C with rotation. Beads were then washed three times with lysis buffer and eluted at 65 °C in 1x urea sample buffer.

### Western blotting and antibodies

Western blot samples were derived from immunoprecipitation experiments described above, or samples were lysed in NP40 buffer (30 mM HEPES-NaOH, 150 mM NaCl, 0.5% NP40) supplemented with protease inhibitors and benzonase for 30 minutes on ice, and clarified by centrifugation at 21,000 x g. Western blot samples were normalized to protein concentration using Pierce 660 nm Protein Assay reagent (Thermo Fisher, 22660). Next, 2x urea sample buffer was added to the samples, which were then denatured at 65 °C. SDS–PAGE and immunoblotting were performed using the indicated antibodies. Images were captured using a ProteinSimple FluorChem M device.

The following antibodies were used in this study, with specific dilutions varying depending on the experiment: anti-PPAT (rabbit, ProteinTech, 15401-1-AP); anti-NUDT5 (rabbit, ProteinTech, 27004-1-AP); anti-PYCR2 (rabbit, ProteinTech, 17146-1-AP); anti-HPRT (rabbit, ProteinTech,15059-1-AP); anti-vinculin (rabbit, CST, 4650); anti-FLAG (mouse, Clone M2; Sigma, F1804); anti-DYKDDDDK Tag (rabbit, CST, 1473) anti-NRF2 (rabbit, CST, D1Z9C); anti-NEDD8 (rabbit, CST, 19E3); anti-α-tubulin (mouse, DM1A, Calbiochem, CP06); anti-V5 (rabbit, CST, D3H8Q); anti-HA (rabbit, CST, C29F4).

### Protein expression and purification

#### PPAT

PPAT^V5-TwinStrep^ was lentivirally expressed in Expi293F human cells in which NUDT5 was bulk-depleted using sgRNAs a described above. Protein preparations were performed from between 2-6 liters of culture harvested at ∼8x10^6^ cells/mL. Purification buffers were extensively vacuum degassed and sparged with nitrogen gas prior to being cooled on ice. All purification steps were carried out on ice or at 4 °C. Cell pellets were resuspended in lysis buffer (50 mM HEPES-NaOH pH 7.5, 150 mM NaCl, 1 mM DTT) supplemented with EDTA-free protease inhibitor cocktail and benzonase. Cells were lysed by brief sonication and clarified by centrifugation. The supernatant was treated with BioLock reagent (1 mL/40 mL lysate; IBA, 2-0205-050) for 20 minutes then incubated with Streptactin XT 4Flow resin for 90 minutes with gentle agitation. Resin was transferred into a gravity column and washed with 10-column volumes (CV) of lysis buffer followed by 20-CVs of lysis buffer supplemented with 10 mM MgCl_2_ and 10 mM ATP heated to 37 °C prior to application to remove chaperone contamination, and an additional 10-CVs of cold lysis buffer. Protein was eluted in lysis buffer containing 50 mM biotin. The eluate was concentrated (Amicon, UFC9050) and buffer exchanged using a 0.5 mL Zeba desalting column into lysis buffer without biotin (7 kDa MWCO; ThermoFisher Scientific, 89882), flash-frozen in small aliquots, and stored at -80 °C for future use.

#### NUDT5

His-SUMO-TEV-NUDT5 wild-type and mutants were expressed in LOBSTR-BL21(DE3)-RIL cells (Vector Laboratories, NC1789768) induced with 0.5 mM IPTG for ∼16 hours at 18 °C. Cell pellets were resuspended in lysis buffer (50 mM Tris-HCl pH 8, 500 mM NaCl, 10 mM imidazole) supplemented with EDTA-free protease inhibitor cocktail and phenylmethylsulfonyl fluoride (PMSF), benzonase, and lysozyme. Cells were lysed via sonication and the supernatant clarified by centrifugation. Supernatant was applied to a 5 mL HisTrap crude FF column (Cytiva, 17525501) via syringe pump. The column was washed with 50 mL of lysis buffer followed by 25 mL of wash buffer (50 mM Tris-HCl pH 8, 500 mM NaCl, 30 mM imidazole) and eluted in 20 mL of elution buffer (50 mM Tris-HCl pH 8, 200 mM NaCl, 300 mM imidazole). Eluate was incubated with TEV protease (1:50 w/w; produced in house by MacroLab, UC Berkeley) and dialyzed overnight against 4L of buffer (50 mM Tris-HCl pH 8.0, 200 mM NaCl, 2 mM BME). The cleaved product was filtered over a HisTrap column, the flow-through concentrated, and applied to a HiLoad Superdex 75 preparative SEC column (Cytiva, 28989333) equilibrated in storage buffer (25 mM HEPES-NaOH pH 7.5, 150 mM NaCl, 1 mM DTT) using an AKTA pure 25 FPLC (Cytiva).

#### PPAT-NUDT5 for cryo-EM (AMP complex)

PPAT^V5-TwinStrep^ and 3xFLAG-NUDT5^Y74F^ were co-expressed lentivirally in 4 liters of wild-type Expi293F cell culture grown in Expi293 media supplemented with 100 μM adenine and harvested at ∼8x10^6^ cells/mL. All purification steps were performed on ice or at 4 °C in purification buffer (50 mM HEPES-NaOH pH 7.5, 150 mM NaCl, 2 mM AMP, 1 mM DTT). The first strep-tag immunoprecipitation step was performed as described above to purify PPAT, omitting the MgCl_2_/ATP wash. Eluate from the Streptactin XT resin was rebound to 1 mL of FLAG-M2 agarose resin for two hours in purification buffer without DTT, washed three times, and eluted three times in a total of 6 mL of purification buffer supplemented with 1 mg/mL 3xFLAG peptide (Sigma Aldrich, F4799). The FLAG-M2 eluate was concentrated in a 100 kDa MWCO concentrator (Amicon) and applied to a Superose6 10/300 increase SEC column (Cytiva, 29091596) equilibrated in SEC buffer (25 mM HEPES-NaOH pH 7.5, 150 mM NaCl, 2 mM AMP, 1 mM DTT). The complex was concentrated to ∼9 mg/mL and immediately used for cryo-EM grid preparation.

#### PPAT-NUDT5 for cryo-EM (6-meTIMP and 6-benzylTIMP complexes)

PPAT^V5-TwinStrep^ and bacterially expressed NUDT5^Y74F^ were purified separately as described above. In this case, we used NUDT5^Y74F^ due to observations that it forms a more stable complex with PPAT than the wild-type protein. NUDT5 (75 μM) was incubated with PPAT (15 μM) directly after elution of PPAT from Streptactin resin for one hour in 2 mL of Streptactin elution buffer (50 mM HEPES-NaOH pH 7.5, 150 mM NaCl, 1 mM DTT, 50 mM biotin) supplemented with 200 μM 6-meTIMP (Jena Biosciences, NU-1226) or 6-benzylTIMP (WuXi). The protein mixture was concentrated to 350 μL and the concentration of nucleotide adjusted to 1 mM. After an additional 1-hour incubation, the complex was applied to a Superose6 10/300 increase SEC column (Cytiva) equilibrated in SEC buffer (25 mM HEPES-NaOH pH 7.5, 150 mM NaCl, 50 μM 6-meTIMP or 200 μM 6-benzylTIMP, 1 mM DTT). Fractions containing the PPAT-NUDT5 complex were concentrated to ∼15 mg/mL. The complex was diluted to 10 mg/mL and a final concentration of 2 mM 6-meTIMP/6-benzylTIMP in cryo-EM buffer (25 mM HEPES-NaOH pH 7.5, 150 mM NaCl, 1 mM DTT) and used immediately for grid preparation.

### Cryo-EM sample preparation, data collection, and analysis

Cryo-EM samples were mixed with a final concentration of 0.02% (w/v) fluorinated octylmaltoside (Anatrace, O310F) immediately before cryo-freezing to prevent protein denaturation at the air–water interface. Then, 2.6 μL of the sample was applied to a glow-discharged 300-mesh Quantifoil R1.2/1.3 grid and incubated for 15 s before being blotted and plunge-vitrified in liquid ethane cool-protected by liquid nitrogen. Grid freezing was performed using a Mark IV Vitrobot (Thermo Fisher Scientific) system operating at 12 °C and 100% humidity.

Cryo-EM data were collected using a 300 kV Titan Krios G3i microscope (Thermo Fisher Scientific) equipped with a BIO Quantum energy filter (slit width 20 eV). Data were collected using SerialEM software at a nominal magnification of 105,000x with a pixel size of 0.424 Å per pixel. Movies were recorded using a 6k x 4k Gatan K3 Direct Electron Detector operating in super-resolution CDS mode. Each movie was composed of 40 subframes with a total dose of 50 e-/A^2^, resulting in a dose rate of 1.25 e-/A^2^. Data processing, including motion correction, CTF estimation, particle picking, 2D class averaging, and 3D refinement, was performed using cryoSPARC v.4.3 workflow ^77^, using mostly default settings. All movies were 2x binned and patch motion corrected. After particle picking and several iterations of 2D class averaging, the initial 3D volume was calculated using several rounds of ab initio 3D reconstruction. Selected particles were then used for non-uniform 3D refinement ^78^. Default B-factor sharpening or local filtering was used to generate the final sharpened maps.

### Model building

The initial model for the PPAT-NUDT5 complex was generated using AlphaFold3 ^48^ by specifying four chains of NUDT5 (UniProt ID: Q9UKK9), four chains of PPAT (UniProt ID: Q06203), and 8 molecules of AMP. Coordinates for 4Fe-4S clusters were added by aligning *B. subtilis* PPAT (PDB ID: 1GPH) to the AlphaFold model. The model was fit into the high-resolution cryo-EM density map in ChimeraX (v1.9) ^79^, and model regions without adequate density in the unsharpened map were deleted. Models were iteratively refined using a combination of ISOLDE (v1.2) ^80^ in ChimeraX (v1.9), Coot (v0.9.4.1) ^81^, and phenix.real_space_refine and phenix.validation_cryoem in Phenix (v1.21.2-5419) ^82^. Models from the AMP-bound datasets were used as initial models for the additional datasets. The Grade2 server was used to generate ligand structure and restraints for the 6-meTIMP and 6-benzylTIMP (Global Phasing Ltd).

### PPAT activity assay

PPAT activity assays monitoring glutamate formation were performed in 96-well PCR plates at a final volume of 20 μL, and all reaction components were diluted in PPAT reaction buffer (50 mM HEPES-NaOH pH 7.5, 150 mM NaCl, 5 mM MgCl_2_, 0.1 mg/mL BSA, 1 mM DTT). First, 5 µL of nucleotide dilution was added to each well, followed by 5 µL of buffer or NUDT5 ([C]_f_ = 10 µM), and then 5 μL of PPAT ([C]_f_ = 50 nM). Reaction components were incubated at room temperature for 30 minutes, then initiated by addition of 5 µL of 4x substrate mixture containing PRPP ([C]_f_ = 1mM; Sigma Aldrich) and glutamine ([C]_f_ = 4mM; Sigma Aldrich). Some assays were performed with 0.25 mM PRPP to better measure competitive inhibitor potency, indicated in the figure legends. Reactions proceeded for 8 minutes at 37 °C in a thermocycler and were quenched at 98 °C for 2 minutes. Reaction kinetics were observed to be linear for 12.5 mins for reactions at both 0.25 and 1 mM PRPP. Reactions were diluted 1:100 in 10 mM HEPES-NaOH pH 7.5, and 20 µL of dilution was transferred to a white 384-well microplate (Corning, 3752). 5 μL of Glutamate-Glo reagent (Promega, J7021) was added to each well, incubated for two hours, and luminescence signal was measured on a Perkin Elmer EnVision microplate reader.

Raw data were fit to a four-parameter logistic equation, Y = Ymin + (Ymax − Ymin) / [1 + (X / 10^logKi)^n] in Python (v3.11.8) using the scipy.optimize.curve_fit function and fits were manually inspected. Inhibition curves that reached saturation were baseline normalized using both Ymin and Ymax values, whereas conditions that did not reach saturation were normalized using only Ymax. Normalized data were plotted in GraphPad prism (v10) and fit using the “[Inhibitor] vs. response (three parameters)” equation with the bottom value fixed at 0 and hill coefficient fixed at 1, which was the approximate hill coefficient for inhibition curves in the presence of wild-type NUDT5. In general, conditions without NUDT5 exhibited n>1, but we opted to constrain this value in order to not over-fit wild-type NUDT5 inhibition curves for direct comparison. The reported Ki is the mean ± SEM from n = 3 independent experiments.

### Mass spectrometry metabolomics

#### Isotope tracing experiments

Cells were seeded in 6 cm dishes in DMEM with 10% dialyzed FBS (dFBS) and 2 mM L-Glutamine. For experiments with hypoxanthine, 20 μM was included at plating. For experiments with 6-MP, 20 μM of drug was added ∼36 hours after plating and ∼12 hours before addition of isotopes for tracing. Around 48 hours after plating, media was exchanged by washing plates 1 time in a minimal media (DMEM with dFBS) before adding media containing either [^15^N-amide]-L-glutamine (2 mM; Cambridge Isotope Labs, NLM-557) or unlabeled L-glutamine (2 mM) with fresh hypoxanthine or 6-MP (20 µM) supplemented for the described treatments. After labeling for 3 hours, cellular metabolism was quenched by aspirating media, snap freezing plates in liquid nitrogen, and storing at -80 °C until extraction. Plates were harvested in parallel accounting for cell number for proactive normalization during sample processing. Intracellular metabolites were extracted by adding extraction solvent (40% acetonitrile, 40% methanol, 20% water, and 0.1 M formic acid with 1 μM ^13^C_6_-glucose-6-phosphate and 10 μM ^13^C_1_-fructose-1,6-bisphosphate internal isotopic standards) to each dish (7.5x10^6^ cells/mL extraction solvent) and incubated at 4 °C for 10 min. Cell extracts were scraped, neutralized with ammonium bicarbonate (0.1 M final), and stored at -80 °C until analysis. Prior to injection, cell extracts were centrifuged at 17,000 x g for 10 minutes at 4 °C, and supernatant was used for analysis.

Metabolites from cellular extracts were injected (10 μL) and separated using the SeQuant ZIC-pHILIC column (5 mm polymeric sorbent, 150 × 2.1 mm; Millipore-Sigma, 1.50460.0001) using an Agilent 1260 Infinity HPLC. Autosampler employed a 10-second needle wash between samples (1:1:1 isopropanol, acetonitrile, water) and was maintained at 10°C. Chromatographic separation was performed using a 30-minute linear gradient starting and 10% ammonium acetate (20 mM, pH 9.3) with medronic acid (5 μM; Agilent, 5191-4506) and 90% acetonitrile, terminating at 30% acetonitrile. Flow rate and column temperature were maintained at 200 μL per min and 15°C, respectively. A triple quadrupole mass spectrometer (Agilent 6430 QQQ) equipped with electrospray ionization was coupled to the HPLC system to perform targeted metabolomics via dynamic multiple reaction monitoring (dMRM). The dMRM method utilized both positive and negative ionization mode, where metabolite m/z and retention times optimized and validated using an in-house metabolite library. QQQ source parameters were held constant at: gas, 350C at 11 L/minute; nebulizer, 25 psi; capillary, 3000V (both negative and positive). Agilent MassHunter (version 10.1) was used for data acquisition, and Skyline (MacCoss Lab version 24.1.0.214) ^83^ was used for analysis.

Each independent experiment contained three biological replicates per sample per treatment group. To quantify relative abundance of total metabolite pools in unlabeled samples, the total peak area for all measured isotopologs were normalized to the isotopic standard with the closest retention time. For de novo purine synthesis rates in [^15^N-amine]-glutamine labeled samples, isotopic enrichment was calculated using the raw peak areas of the fully labeled isotopolog relative to the sum of all biologically relevant isotopologs. For AMP and IMP, relevant isotopologs included M+0 and M+2, where M+2 was deemed fully labeled. For GMP, relevant isotopologs included M+0, M+2, and M+3, where M+3 was deemed fully labeled.

#### Thiopurine metabolite abundance measurements

Intracellular levels of 6-TIMP, and 6-meTIMP were determined via LC-MS/MS. Cells were washed with cold PBS prior to the addition of a 40/40/20 mixture of LC-MS grade acetonitrile/methanol/water +0.1% formic acid for intracellular metabolite extraction. Cells were incubated for 30 mins in -20°C, then centrifuged at 14,000 RPM for 10 mins to pellet insoluble material. Isotope labeled ^13^C-^15^N amino acids (Cambridge Isotope Laboratories, MSK-A2-1.2) were also spiked into each sample as internal standards to monitor analytical conditions. Supernatant was transferred to LC-MS vials for metabolomics analysis where 2uL of sample was used for analysis.

Untargeted metabolomics was conducted on a Vanquish UHPLC system coupled with an Orbitrap Exploris^TM^ 240 mass spectrometer (Thermo Fisher Scientific). Polar metabolites were separated on a XBridge Amide Column (2.1 mm ID × 100 mm, particle size 3.5 μm; Waters,186004860) which was consistently housed at 40°C. Mobile phases were prepared as following: (A) 95/5 (v/v) water/acetonitrile with 10 ammonium hydroxide and 10 ammonium acetate; (B) 5/95 (v/v) water/acetonitrile with 10 mM ammonium hydroxide and 10 mM ammonium acetate. The mobile phases were delivered at a flow rate of 0.15 mL/min for a 25 min run with the following stepwise gradient for solvent B: 0 min, 85% B; 2.5 min, 70% B; 7 min, 55% B; 16 min, 35 % B; 16.1-18 min, 25 % B; 18-25 min, 85 % B. The electrospray ionization source (ESI) was operated in positive mode. The ion spray voltage was set at 4 kV with the ion transfer tube temperature set at 350°C, vaporizer temp 325°C, sheath gas 35 arbitrary units (au), aux gas 5 au and sweep gas 1 au. Full MS scans of 1 ms were performed at 90,000 units resolution, with a scan range of 60-900 m/z. A pooled quality control (pQC) sample followed by a blank (mobile phase solution) injection was performed among every ten injections of biological samples to monitor instrument performance. The pQC sample were also utilized for the top 10 MS/MS analyses with dynamic exclusion during the analysis for compound identification. Peaks were integrated using the Quan Browser module within Xcalibur and were normalized to protein content for each sample.

### Mass spectrometry proteomics

#### Sample processing

FLAG-M2 beads from immunoprecipitations were diluted in TNE (50 mM Tris pH 8.0, 100 mM NaCl, 1 mM EDTA) buffer. RapiGest SF reagent (Waters, 186008090) was added to the mix to a final concentration of 0.1%, and the samples were boiled for 5 min. TCEP was added to 1 mM (final concentration) and the samples were incubated at 37 °C for 30 min. Subsequently, the samples were carboxymethylated with 0.5 mg ml−1 of iodoacetamide for 30 min at 37 °C followed by neutralization with 2 mM TCEP (final concentration). The proteins samples were then digested with trypsin (trypsin:protein ratio, 1:50) overnight at 37 °C. RapiGest was degraded and removed by treating the samples with 250 mM HCl at 37 °C for 1 h followed by centrifugation at 14,000 rpm for 30 min at 4 °C. The soluble fraction was then added to a new tube and the peptides were extracted and desalted using C18 desalting columns (Thermo-Fisher Scientific, PI-87782). Peptides were quantified using BCA assay and a total of 1 μg of peptides were injected for LC-MS analysis.

#### LC-MS analysis

Trypsin-digested peptides were analyzed by ultra-high-pressure liquid chromatography (UPLC) coupled with tandem mass spectroscopy (LC–MS/MS) using nano-spray ionization. The nanospray ionization experiments were performed using a TimsTOF 2 pro hybrid mass spectrometer (Bruker) interfaced with nanoscale reversed-phase UPLC (EVOSEP ONE). The Evosep method of 30 samples per day was performed using a 10 cm × 150 μm reversed-phase column packed with 1.5 μm C18-beads (PepSep, Bruker) at 58 °C. The analytical columns were connected with a fused silica ID emitter (10 μm inner diameter, Bruker Daltonics) inside a nanoelectrospray ion source (captive spray source, Bruker). The mobile phases comprised 0.1% formic acid as solution A and 0.1% formic acid/99.9% acetonitrile as solution B. The MS settings for the TimsTOF Pro 2 were as follows: the dia-PASEF method for proteomics. The values for mobility-dependent collision energy were set to 10 eV. No merging of TIMS scans was performed. The ion mobility (IM) was set between 0.85 (1/k0) and 1.3 (1/k0) with a ramp time of 100 ms. Each method includes one IM window per dia-PASEF scan with variable isolation window at 20 amu segments; 34 PASEF MS/MS scans were triggered per cycle (1.38 s) with a maximum of 7 precursors per mobilogram. Precursor ions in an m/z range of between 100 and 1,700 with charge states ≥3+ and ≤8+ were selected for fragmentation. Protein identification, label-free quantification and statistical analysis were performed using default settings in Spectronaut 18.0 (Biognosys).

For the PPAT^TwinStrep^-NUDT5^3xFLAG^ sequential IP and NUDT5^3xFLAG^ IP comparison, DIA protein quantity for candidate hits from both experiments were normalized to that measured for NUDT5 bait. The resulting gene list was compared to the CRAPome database of common mass spectrometry contaminants, and hits that were found in >50% of database experiments were removed from the analysis.

### Multiple sequence alignments (MSAs)

MSA’s were performed using Clustal Omega ^84^. The UniProt sequence identifiers are as follows (species, NUDT5 accession, PPAT accession): human (Homo sapiens, Q9UKK9, Q06203); primate (Macaca mulatta, A0A1D5QNV6, F7GPV9); mouse (Mus musculus, Q9JKX6, Q8CIH9); rat (Rattus norvegicus, Q6AY63, P35433); chicken (Gallus gallus, A0A8V0XJK7, P28173); frog (Xenopus laevis, Q6IND3, A0A974DTF4); fish (Danio rerio, Q6IQ66, A6H8S4).

### Software and programs

All software used is freely/commercially available: FACSDiva (Version 9.0), FlowJo (Version 10.10.0), GraphPad Prism (Version 10), SerialEM (Version 4.1), AlphaFold 3, Coot (Version 0.9.8.92), ChimeraX (Version 1.8), PyMOL (Version 2.5.5), PHENIX (Version 1.21.1-5286), cryoSPARC (Version 4.3), Isolde (Version 1.2), Spectronaut (Version 18.0), Agilent MassHunter (Version 10.1), Skyline (MacCoss Lab version 24.1.0.214), Xcalibur (Version 4.3). Python (v3.10), SciPy library (v1.11), Matplotlib (v3.7).

## Data availability

Structural models of PPAT-NUDT5-nucleotide complexes have been deposited to the Worldwide Protein Data Bank (wwPBD) and Electron Microscopy Data Bank (EMDB) with the following accession codes: AMP complex (PDB: 9Q0M; EMDB: EMD-72099), 6-meTIMP complex (PDB: 9Q0N; EMDB: EMD-72100), 6-benzylTIMP complex (PDB: 9Q0O; EMDB: EMD-72101).

Numerical and immunoblot source data are provided as supplementary data with this manuscript. Unprocessed MS data files from proteomics experiments were deposited to the ProteomeXchange Consortium via the PRIDE partner repository (https://www.ebi.ac.uk/pride/archive) under dataset identifier XXXX. There are no restrictions on data availability.

## Acknowledgements

We thank all of the members of our laboratory for enthusiastic support and many suggestions for this work; K. Birsoy (Rockefeller University) for sharing the human metabolism-focused sgRNA library; QB3 Genomics, UC Berkeley, Berkeley, CA, for Illumina sequencing; D. Toso and the members of the Cal-Cryo cryo-EM facility at QB3-Berkeley for assistance on cryo-EM data collection; the UCB Cell Culture Facility which is supported by The University of California Berkeley (RRID is SCR_017924); and M. Ghassemian and the members of the UCSD Biomolecular and Proteomics Mass Spectrometry facility for assistance in running the MS samples. We thank the Molecular Therapeutics Initiative at UCB for chemistry advice and support with global metabolomics measurements. F.C. was supported by the Molecular Therapeutics Initiative. S.R.W. was supported by a fellowship from the American Cancer Society (PF-24-1257623-01-TBE). H. R. is a Howard Hughes Medical Institute Fellow of The Jane Coffin Childs Memorial Fund. This work was supported by the Howard Hughes Medical Institute (MR) and R35 GM152114 (DT). M.R. is an Investigator of the Howard Hughes Medical Institute.

## Contributions

S.R.W. conceived of and performed most of the experiments and analyses including CRISPR screening, cryo-EM analysis, cell culture experiments, and *in vitro* reconstitutions. M.M.K. and F.C. performed metabolomics-based experiments. H.R. performed immunoprecipitations and cell growth assays. Z.Y. collected and processed cryo-EM data. A.S.G. supported biochemical analysis of the PPAT-NUDT5 complex. M.R. and D.V.T. helped to plan and interpret experiments. S.R.W. and M.R. wrote the manuscript with input from all authors.

---

**Extended Data Figure 1.**
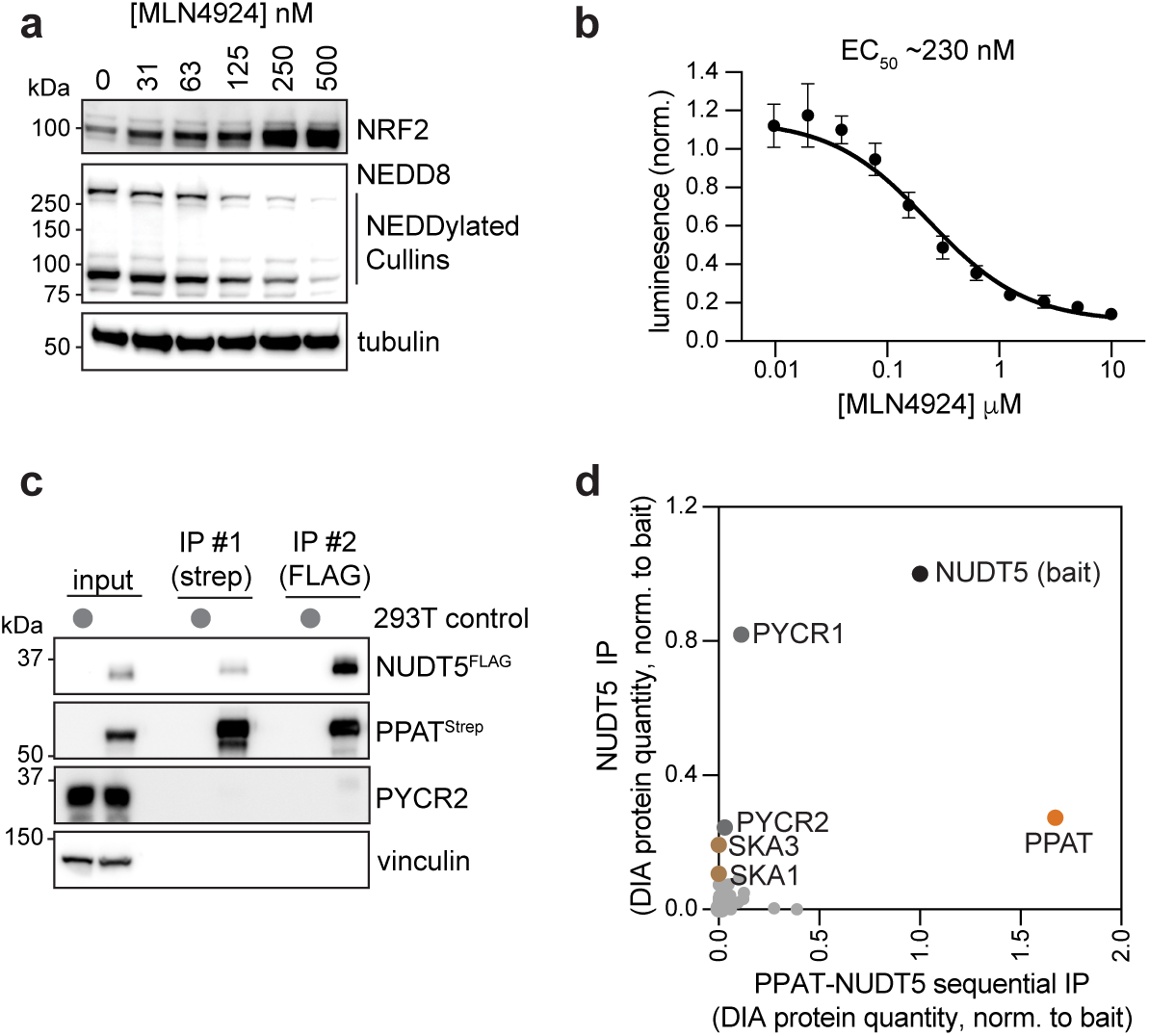
Identification of PPAT-NUDT5 complex. **a.** Western blot of U2OS cells treated with indicated amounts of MLN4924 for 16 hours. Data are representative of two independent experiments. **b.** CellTiter Glo cell viability assay of U2OS cells treated with MLN4924 for 72 hours. Data are the mean and error bars are SEM of n=3 biological replicates from independent experiments. **c.** Western blot of sequential immunoprecipitation from HEK293T cells expressing PPAT^Strep^ and NUDT5^3xFLAG^ **d.** Mass spectrometry of PPAT^Strep^-NUDT5^3xFLAG^ sequential immunoprecipitation compared to NUDT5^3xFLAG^ -only immunoprecipitation and normalized to NUDT5 bait protein abundance. Data show the mean of n=3 technical replicates for each condition.

**Extended Data Figure 2.**
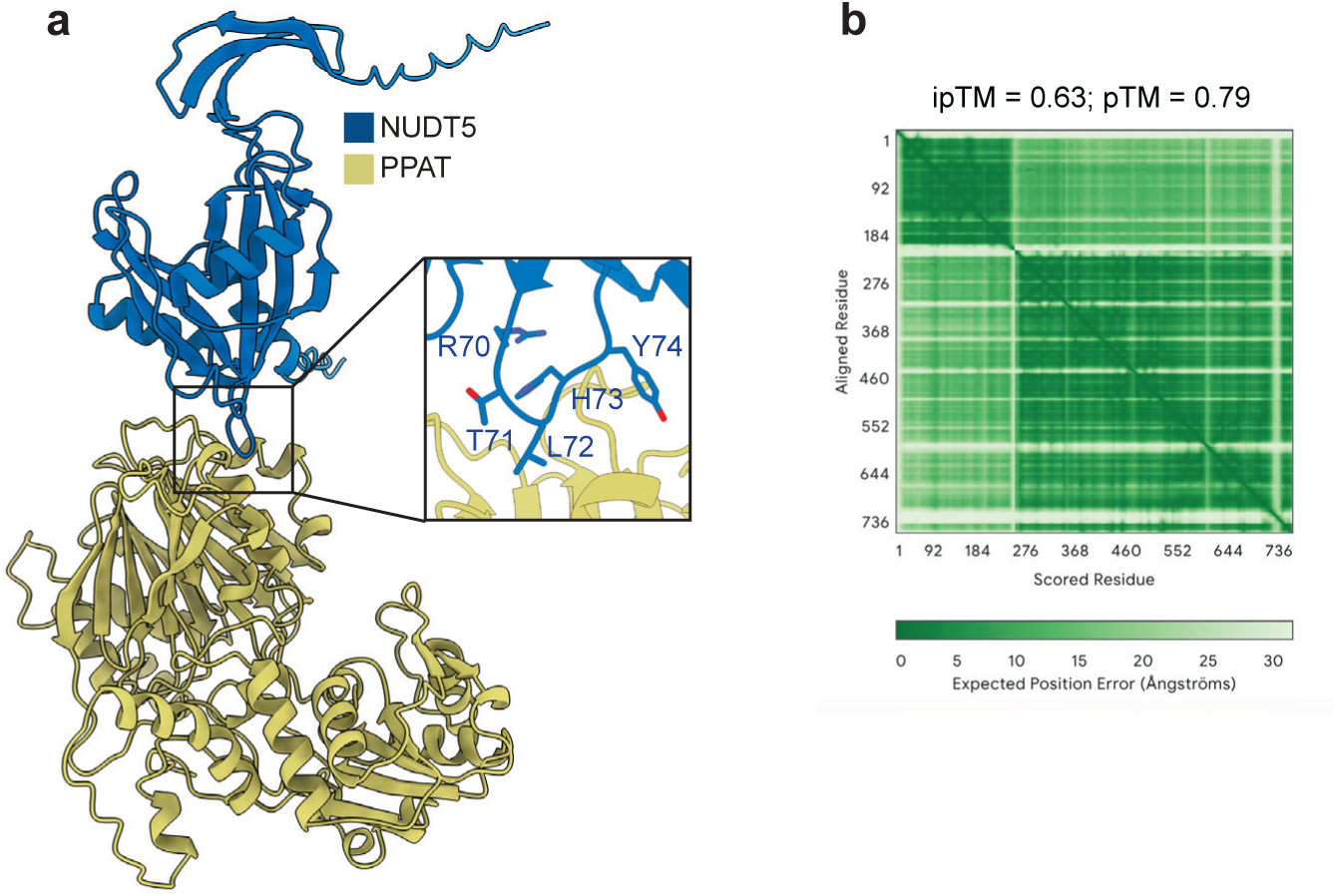
AlphaFold modeling of PPAT-NUDT5. **a.** AlphaFold3 model of PPAT-NUDT5 monomer interaction. **b.** Contact map for structure prediction shown in panel a.

**Extended Data Figure 3.**
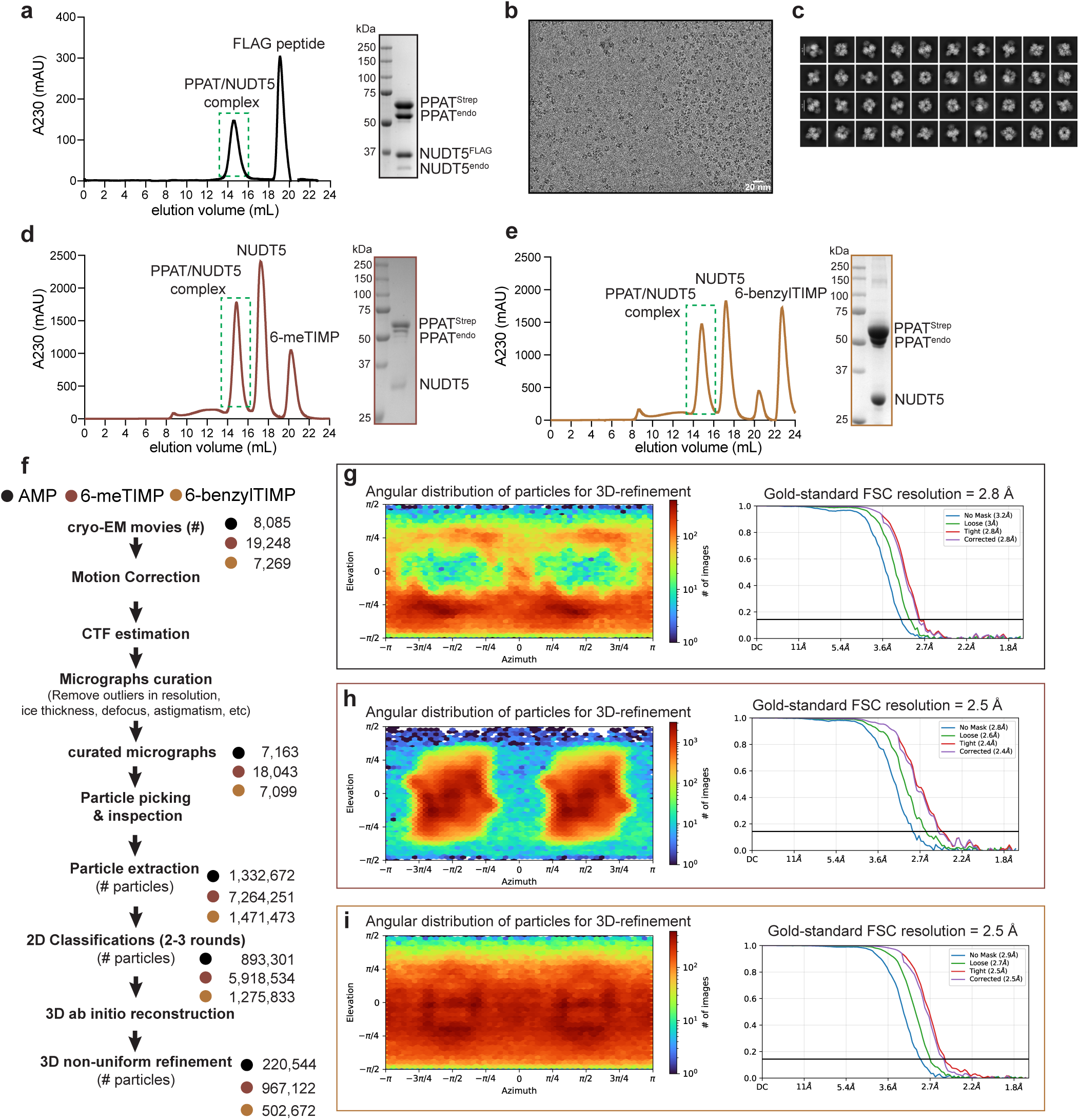
Cryo-EM sample preparation and workflow. **a.** Size exclusion chromatography (SEC) trace (left) and Coomassie-stained gel of PPAT-NUDT5-AMP complex (right) used for cryo-EM analysis. **b.** Motion-corrected and denoised micrograph of PPAT-NUDT5-AMP complex. **c.** Representative 2D-class averages for PPAT-NUDT5-AMP complex. Micrographs and 2D class averages from the 6-meTIMP and 6-benzylTIMP datasets were highly similar to that of the AMP-bound dataset. **d, e.** SEC and Coomassie-stained gels of PPAT-NUDT5 complexes formed in the presence of **d.** 6-meTIMP and **e.** 6-benzylTIMP. **f.** Cryo-EM data processing particle flow for the three nucleotide-bound PPAT-NUDT5 structures. All processing steps were performed in CryoSparc. **g-i.** Angular distribution of particles and FSC curves from refinements of PPAT-NUDT5-purine complexes with **g.** AMP, **h.** 6-meTIMP and **i.** 6-benzylTIMP.

**Extended Data Figure 4.**
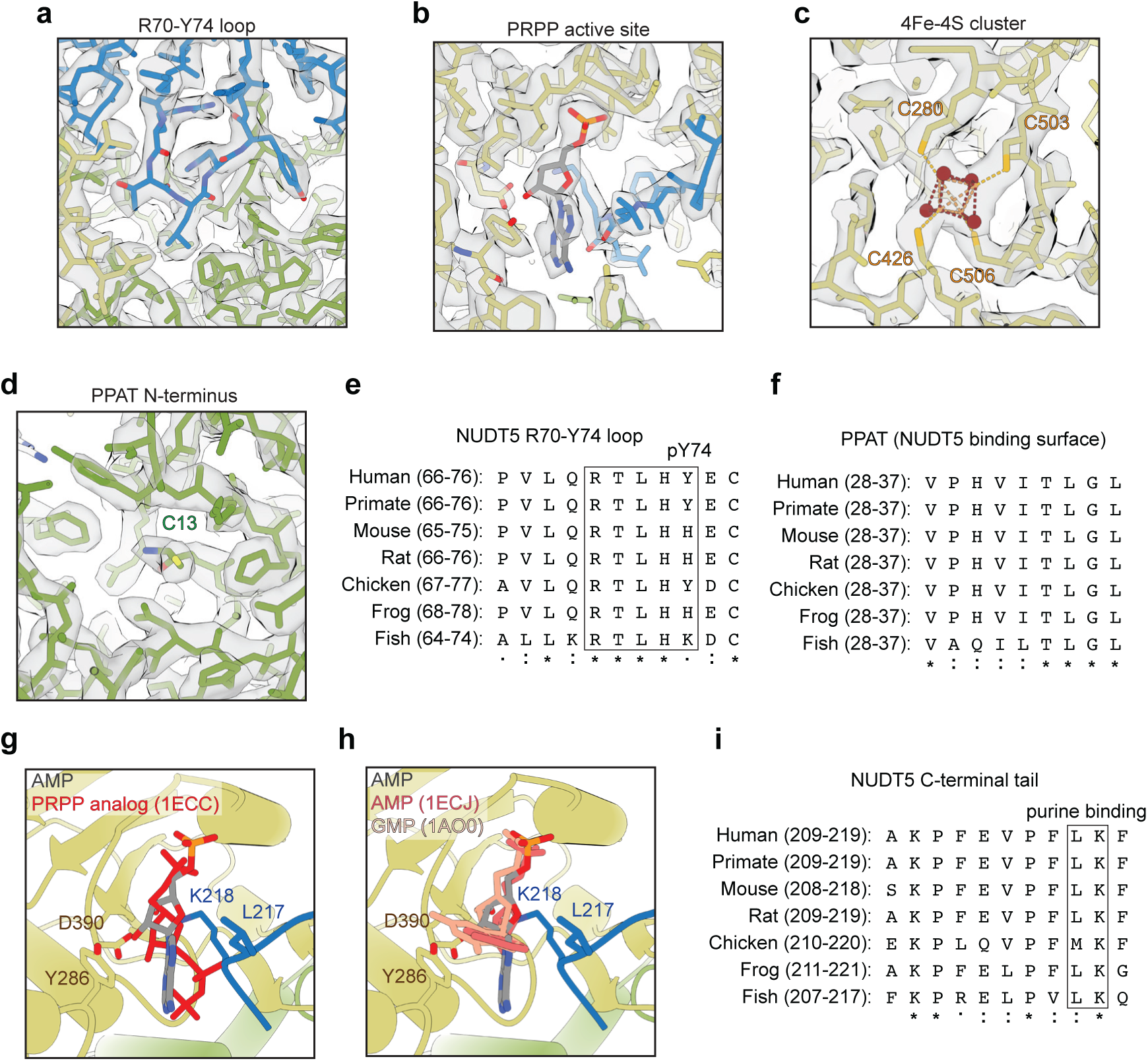
Structural and sequence analysis of PPAT-NUDT5-AMP complex. a-d. Example cryo-EM density from the PPAT-NUDT5-AMP map with fit molecular model highlighting: **a.** the R70-Y74 interface **b.** AMP-bound at the molecular glue interface/PRPP active site. **c.** the 4Fe-4S cluster bound by PPAT and **d.** the processed N-terminus of PPAT. **e, f.** Multiple sequence alignment (MSA) of **e.** NUDT5 R70-Y74 loop region and **f.** A NUDT5-binding region of PPAT. **g, h.** Alignment of PPAT protomers from the human PPAT-NUDT5-AMP complex and **g.** E. *coli* PPAT bound to a non-hydrolysable PRPP analog (PDB: 1ECC), **h.** E. *coli* PPAT bound to AMP (PDB: 1ECJ) and B. *subtilis* PPAT bound to GMP (PDB: 1AO0). Only the nucleotides are shown from the bacterial structures for clarity **i.** MSA of NUDT5 C-terminal tail region.

**Extended Data Figure 5.**
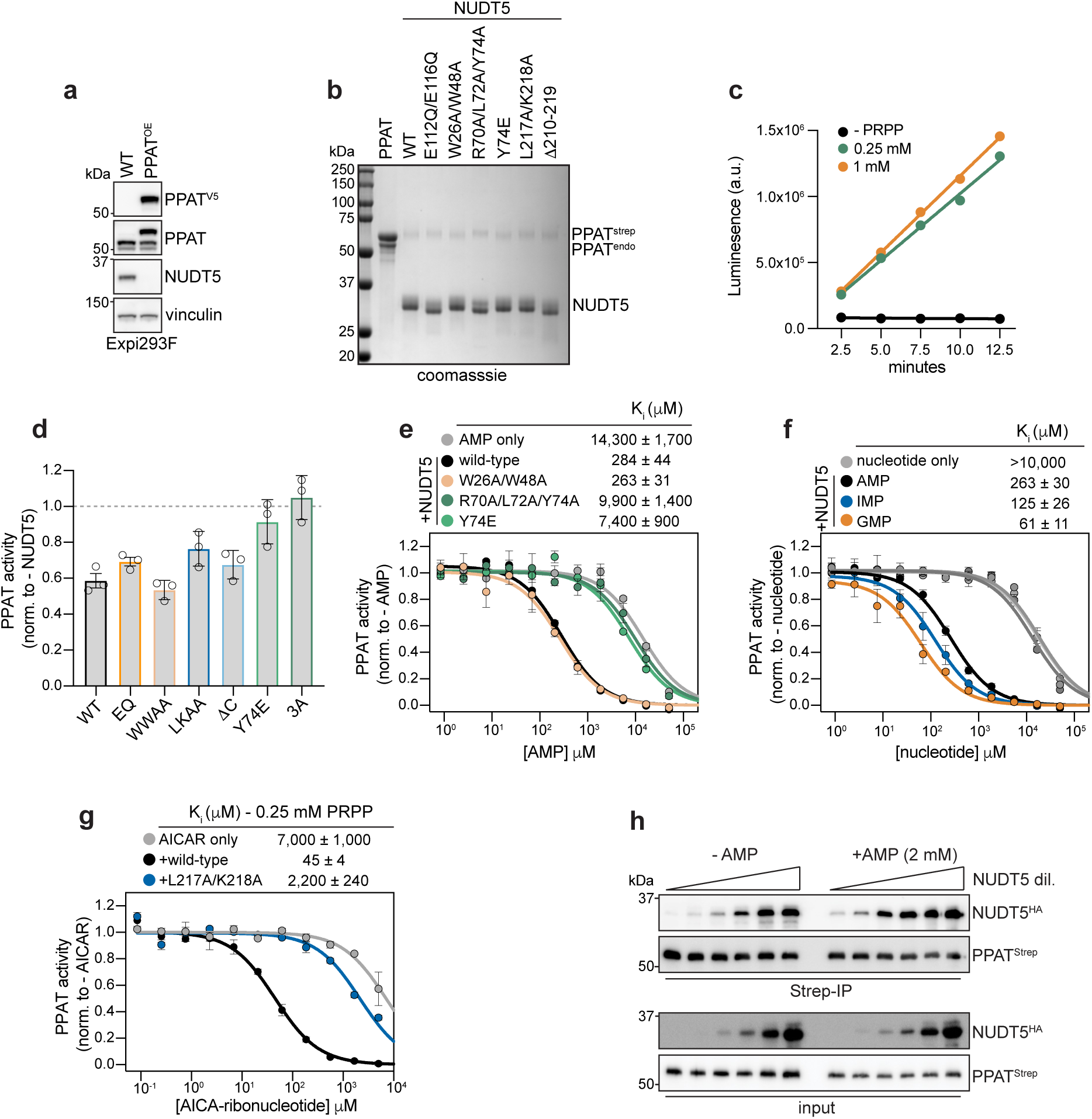
*In vitro* reconstitution of PPAT activity. **a.** Western blot of engineered Expi293F cell line expressing PPAT^V5-TwinStrep^ **b.** Coomassie-stained gel of purified PPAT and NUDT5 proteins. **c.** Time-course PPAT activity assay showing linearity of assays conducted with 0.25 and 1 mM PRPP. Data shown are the mean on n=2 two independent experiments. **d.** PPAT activity with NUDT5 in the absence of AMP conducted with 1 mM PRPP. NUDT5 mutants ‘WWAA’ is W26A/W48A and ‘LKAA’ is L217A/K218A. **e-g.** PPAT activity assay measuring inhibitory effects of **e.** AMP in the presence and absence of NUDT5 wildtype and indicated mutants (1 mM PRPP), **f.** AMP, GMP, and IMP in the presence and absence of wildtype NUDT5 (1 mM PRPP) and **g.** AICA-ribonucleotide in the presence and absence of NUDT5 wildtype and indicated mutants (0.25 mM PRPP). Data in enzyme activity assays in panels d-g show the individual values (d) or mean (e-g) and error bars are SEM from n=3 independent experiments. **h.** Western blot of *in vitro* PPAT^Strep^ immunoprecipitation of recombinant NUDT5 in the presence and absence of AMP. The dilution of NUDT5 in the input is from 10 μM to 2.4 nM (4-fold serial dilution). Data shown are representative from two independent experiments.

**Extended Data Figure 6.**
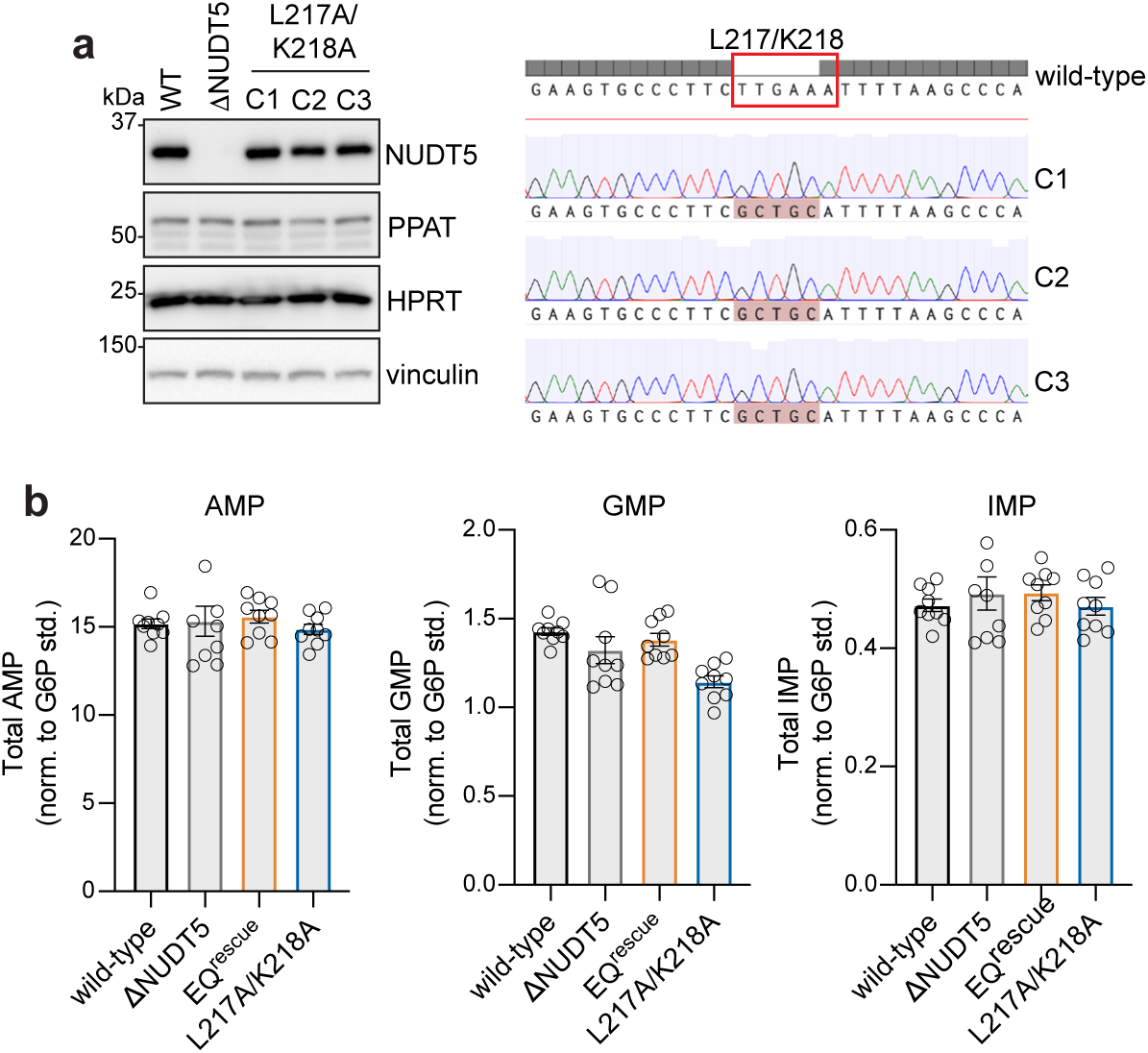
Characterization of endogenous NUDT5 molecular glue mutants. **a.** Left – Western blot of HEK293T cell lines used in this study. Right – aligned sanger sequencing traces of endogenous L217A/K218A clones compared to the wildtype sequence. **b.** Measurements of purine monophosphate levels normalized to a glucose-6’-phosphate internal isotopic standard in HEK293T cells treated with hypoxanthine identically to those used for [^15^N-amide]-glutamine labeling experiments in figure 3. Data points are individual values from n=9 biological replicates from 3 independent experiments and error bars are SEM.

**Extended Data Figure 7.**
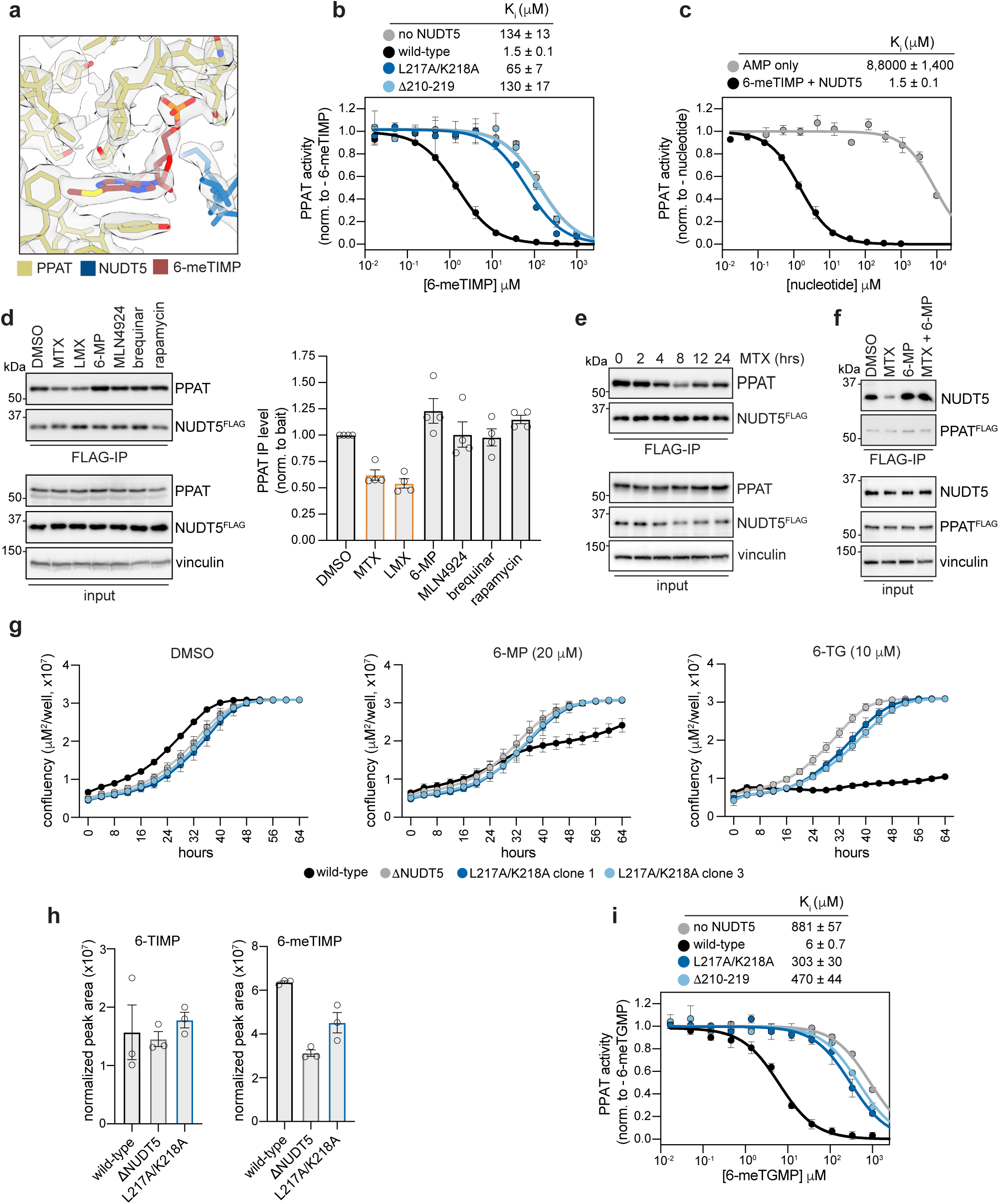
Purine metabolism-targeting chemotherapeutics operate through PPAT-NUDT5. **a.** Example cryo-EM density of the PPAT-NUDT5 6-meTIMP molecular glue interface with model fit. **b,c.** PPAT activity assay measuring inhibitory effects of 6-meTIMP in the presence and absence of wildtype NUDT5 and indicated mutants and **c.** compared to AMP only. **d.** Left – Representative Western blot of immunoprecipitations from endogenous NUDT5^3xFLAG^ HEK293T cells treated with indicated drugs for 16 hours: methotrexate (MTX; 2 µM), lometrexol (LMX; 10 µM), 6-mercaptopurine (6-MP; 50 µM), MLN4924 (1 µM), brequinar (2 µM), and rapamycin (1 µM). Right – quantification of PPAT immunoprecipitation relative to NUDT5^3xFLAG^ bait and normalized to a DMSO-treated control condition. Data are individual values from n=3 biological replicates from independent experiments and error bars are SEM. **e.** Western blot of immunoprecipitations from endogenous NUDT5^3xFLAG^ HEK293T cells treated with MTX (2 µM) for the indicated amounts of time. Similar results were obtained in two independent experiments. **f.** Western blot of endogenous PPAT^3xFLAG^ immunoprecipitations following 16-hour treatment with MTX (2 µM), 6-MP (50 µM), and MTX + 6-MP **g.** Time-resolved microscopy (incucyte) growth assays of wildtype and mutant HEK293T cells treated with the indicated drugs. Data are the mean and error bars are SEM of n=6 biological replicates. **h.** Levels of intracellular 6-TIMP and 6-meTIMP metabolites following 16-hour treatment with 6-MP (20 µM). Data are individual values and error bars are SEM from n=3 biological replicates. **i.** PPAT activity assay measuring inhibitory effects of 6-meTGMP in the presence and absence of wildtype NUDT5 and indicated mutants. Activity data shown in panels b, c and i are the mean and error bars are SEM of n=3 independent experiments.

**Extended Data Figure 8.**
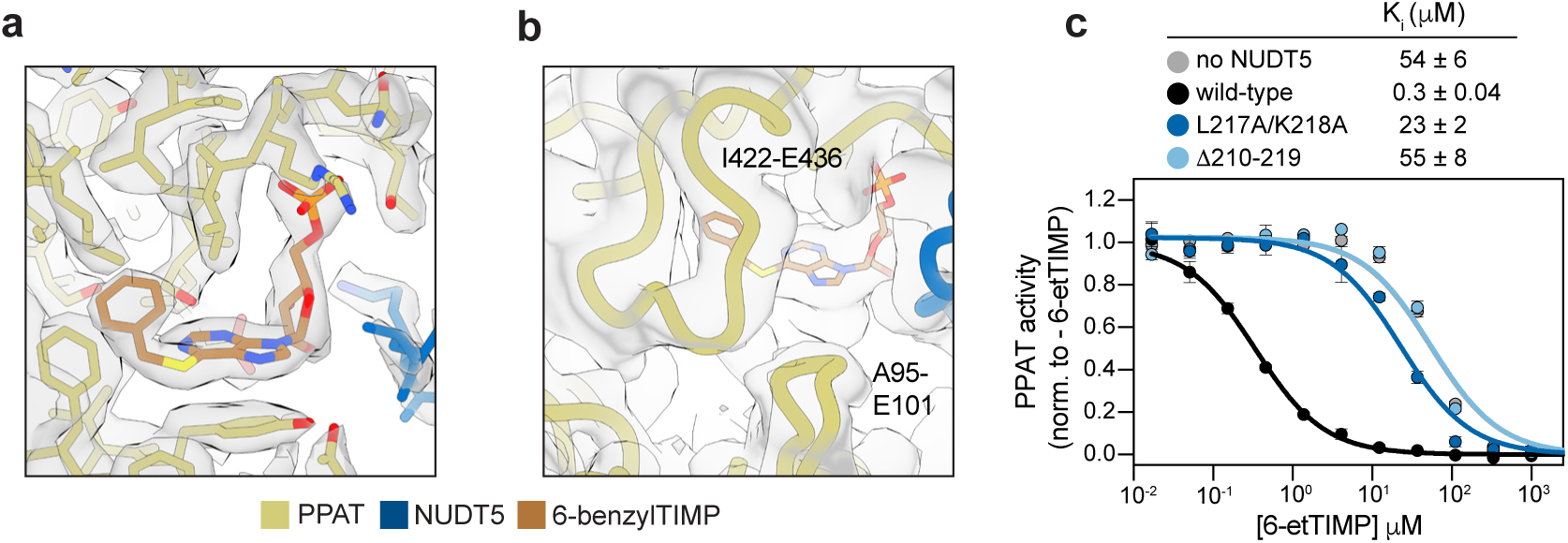
Characterization of thiopurine derivatives. **a.** Example high-resolution cryo-EM density of the PPAT-NUDT5 6-benzylTIMP molecular glue interface with fit model. **b.** I422-E436 and A95-E101 loops of PPAT in the 6-benzylTIMP complex modeled into unsharpened cryo-EM density map and shown at a low contour threshold. **c.** PPAT activity assay measuring inhibitory effects of 6-etTIMP in the presence and absence of wildtype NUDT5 and indicated mutants. Data are the mean and error bars are SEM of n=3 independent experiments.

